# Directional variation and method-specific detection patterns in offshore bat migration: implications for wind farm mitigation

**DOI:** 10.64898/2026.02.06.704313

**Authors:** Sander Lagerveld, Pepijn de Vries, Eldar Rakhimberdiev, Jane Harris, Bart C.A. Noort, Steve C.V. Geelhoed, Frank van Langevelde, Fiona Mathews, Martin J.M. Poot, Julia Karagicheva

**Author notes:** Correspondence*: Sander Lagerveld, Wageningen University & Research, Ankerpark 27, 1781 AG Den Helder, Netherlands.

## Abstract

1. Curtailment of wind farms effectively reduces collision mortality in bats. Implementing this measure in offshore wind farms requires knowledge on the spatiotemporal occurrence and environmental predictors of migration over sea. In bats, such information can be obtained through acoustic monitoring and individual tracking. However, these techniques provide seemingly contradictory insights into migration patterns.
2. We used a Bayesian capture–recapture state-space model to investigate how environmental predictors influence spring departure decisions of Nathusius’ pipistrelle *Pipistrellus nathusii* migrating over the North Sea. The model was applied to both acoustic and tracking data, enabling comparable analyses across methods and incorporating uncertainty in migration dates of tracked bats. Additionally, we examined nightly offshore bat occurrence to further explore differences in movement patterns detected by the two techniques.
3. Wind conditions at 200 m above sea level were identified as key driver of Nathusius’ pipistrelle spring migration. In May–June, most bats migrated from the United Kingdom under westerly and northwesterly tailwinds. Tracked individuals flew in stronger supportive winds than acoustically recorded bats, which were also detected under crosswinds and headwinds. In March–April, acoustic detections occurred mainly during strong southerly winds, suggesting that early-season migrants largely consisted of individuals migrating over the European mainland and drifted northwards onto the North Sea by strong crosswinds.
4. Acoustic detectors primarily recorded bats that landed on offshore platforms, likely because they were unable to cross the North Sea in a single flight due to less favorable wind conditions, or because they departed from more inland locations. In contrast, tracking data mainly represented bats that successfully crossed the North Sea in a non-stop flight under moderate supportive tailwinds.
5. *Synthesis and applications*: Combining observation techniques improves our understanding of bat migration patterns. Additionally, acoustic monitoring can capture migration from different geographic origins. Current mitigation measures for offshore wind farms at the North Sea rely solely on acoustic data, likely overlooking the part of the population that crosses over sea with optimal wind support. Acoustic and tracking data are therefore complementary rather than contradictory, and both methods should be used together when developing mitigation measures.

## 1 Introduction

Worldwide offshore wind farms are expanding rapidly, raising concerns about their potential impact on a suit of species groups. Even non-marine species such as bats can collide with wind turbines during their migration over sea (Ahlén et al., 2009; Cryan and Brown, 2007; Hüppop et al., 2019; Lagerveld et al., 2021, 2014). Significant reductions of the number of bat fatalities have been achieved by the curtailment of wind farms on land, which involves a reduction of the operational time of wind turbines during periods with increased bat activity (Adams et al., 2021; Baerwald et al., 2009; Voigt et al., 2022; Whitby et al.,2024). Accurately predicting bat migration peaks over sea would therefore be of considerable help to develop effective curtailment strategies for offshore wind farms.

A principal area for offshore wind developments is the North Sea, where Nathusius’ pipistrelle *Pipistrellus nathusii* is the most frequently recorded migratory bat (Brabant et al., 2021; Hüppop and Hill, 2016; Lagerveld et al., 2014). The species is particularly vulnerable to collisions with wind turbines (EUROBATS, 2015; Kruszynski et al., 2022) and recent genetic research suggests a population decline (Van Schaik et al., 2025), highlighting the need for conservation measures.

Understanding migration patterns of bats, however, remains challenging due to their elusive and nocturnal lifestyle. To date, most detailed knowledge has come from acoustic monitoring and tracking studies. Acoustic monitoring involves the detection and analysis of bat echolocation calls, and when stationary acoustic detectors are deployed, it enables long-term monitoring of migratory bat activity patterns (Hüppop and Hill, 2016; Ijäs et al., 2017; Johnson et al., 2011; Lagerveld et al., 2021, 2023b; Rydell et al., 2014). Tracking studies on the other hand, provide the most detailed insights into bat migration ecology, including departure and routing decisions (Bach et al., 2022; Dechmann et al., 2017; Hurme et al., 2025; Lagerveld et al., 2024; McGuire et al., 2012; True et al., 2023), and flight behavior during long-distance endurance migratory flights (Lagerveld et al., 2024; O’Mara et al., 2019). For small bat species, such as Nathusius’ pipistrelle, movements over larger distances can currently only be tracked using the Motus Wildlife Tracking System (Mitchell et al., 2025; Taylor et al., 2017), as alternative systems like the Sigfox IoT network (Wild et al., 2023) use tags that are currently too heavy for these small-sized animals.

In some cases, different monitoring methods yield seemingly contradictory insights into the patterns of migration. These discrepancies likely stem from methodological differences in both detection and analysis. For example, tracking data suggest that migration from the east coast of the UK to the European mainland takes place from early May until mid-June (Lagerveld et al., 2024), while acoustic monitoring has shown that bats occur at the southern North Sea from late-March until early June (Lagerveld et al., 2023a). Acoustic data also indicate that sea crossings often span more than one night due to diurnal stopovers at offshore platforms (Lagerveld et al., 2023b), whereas tracking data shows that migratory bats are able to cross the North Sea in a single three to five hour non-stop flight (Lagerveld et al., 2024).

Furthermore, acoustic detections of migratory bats are generally recorded during low wind speeds (Brabant et al., 2021; Cryan and Brown, 2007; Johnson et al., 2011; Lagerveld et al., 2021, 2014; Sjollema et al., 2014), while tracking data and visual observations show that departures more often coincide with moderate or strong winds (Hatch et al., 2013; Lagerveld et al., 2024). Additionally, differences in estimated wind parameters associated with bat departure decisions may arise from methodological limitations inherent to each monitoring technique. For instance, acoustic detectors have a very limited detection range, typically between 25 – 100 m, depending on the species, type of detector and environmental conditions (Adams et al., 2012; Barataud, 2020) -and thus-primarily record low-flying individuals at close range. In contrast, Motus receivers can detect animals flying at altitudes of several kilometers and at distances of 5 - 15 kilometers (Mitchell et al., 2025; Taylor et al., 2017).

However, their capacity to detect low-flying individuals is much more restricted, typically limited to a range of about 2 kilometers (Taylor et al., 2017). Acoustic data thus represents bat activity at low-altitude, potentially underestimating the prevalence of bats migrating at higher altitudes, possibly under different wind conditions. Conversely, low-flying bats may be underrepresented in Motus data.

These discrepancies highlight the need to combine information from data collected with both techniques to better understand the observed offshore occurrence of migratory bats in space and time. Analyzing the two datasets in a comparable way is not only important for the development of curtailment measures to reduce bat fatalities in offshore wind farms, but also for a more comprehensive understanding of bat migration in general.

Between 2018 and 2021, we conducted continuous acoustic monitoring at 13 offshore platforms in the southern North Sea. From 2021 to 2023, we tracked 27 Nathusius’ pipistrelles from the east coast of the UK to the European mainland using the Motus Wildlife Tracking System. With the collected data, we adapted Bayesian capture-recapture models in a state-space formulation (Auger-Méthé et al., 2021; Gimenez et al., 2007; Royle, 2008) to estimate probabilities of bat migratory departures. The flexibility of this type of model allowed its use for both acoustic and tracking data, enabling analysis within a similar analytical framework. Capture-recapture state-space models are also robust to small sample sizes and can account for uncertainty in departure timing. This was particularly useful in the analysis of tracking data that comprised crossings of a limited number of individuals.

By bringing the acoustic and tracking data to the same structure and applying similar -yet separate-analysis, we preserved the distinct characteristics of each monitoring method while enabling direct comparison of outcomes. This approach minimizes methodological biases and enabled us to: 1) examine the species’ spring occurrence throughout the season and the night, 2) identify environmental conditions serving as migratory cues, 3) compare patterns observed across monitoring methods to explain both differences and consistencies in detected occurrence, and 4) recommend mitigation strategies for offshore wind farms.

## 2 MATERIAL & METHODS

### 2.1 Tagging & tracking bats

Bat trapping and tagging were carried out at three sites: Minsmere (52.247° N, 1.619° E), Benacre National Nature Reserve (52.387° N, 1.707° E), and the Landguard Bird Observatory (51.938° N, 1.320° E), over three consecutive years (Figure 1). Fieldwork comprised 14 trapping sessions between 29 March and 15 May 2021 at Minsmere; 10 sessions between 5 April and 10 May 2022 (two at Landguard and eight at Minsmere); and 12 sessions between 29 March and 11 May 2023 (three at Benacre and nine at Minsmere). Trapping sessions lasted an average of 4.5 hours and were conducted under Natural England project licenses 2021-55582-SCI and 2022-63237-SCI.

**Figure 1.**
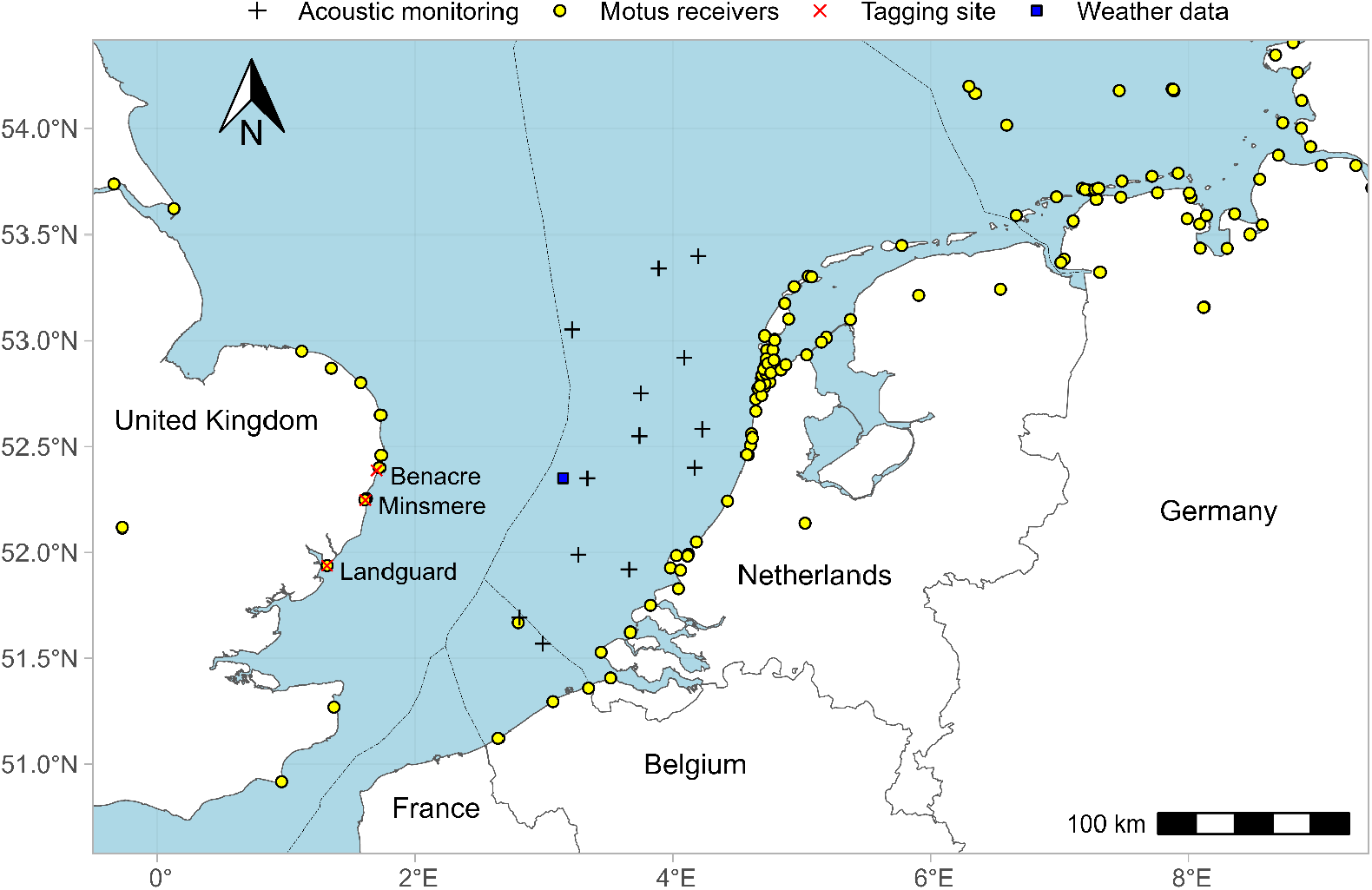
Southern North Sea and bordering countries, including bat tagging sites, offshore acoustic monitoring locations, Motus receiver network and weather data location

We used 0.26 g NTQB2-1 radio-transmitters (LOTEK Wireless, Canada) with burst intervals between 6.7 and 7.3 s and an expected lifespan of 51–61 days. Each transmitter broadcasted a uniquely coded signal at 150.1 MHz, allowing the identification of individual animals. To follow the movements of tagged bats, we used the Motus Wildlife Tracking System (Mitchell et al., 2025; Taylor et al., 2017). In the study area about 125 Motus receiver stations were operational from 2021 – 2023 (Figure 1).

In total, 122 bats were tagged, comprising 18 males and 104 females. The number of individuals tagged differed across years, with 20 in 2021, 24 in 2022, and 78 in 2023. Tagging also varied seasonally, with 2 bats tagged in March, 25 in April, and 95 in May. One individual was predated and two tags were found detached. The remaining 119 bats were all detected in the UK and 27 individuals were subsequently recorded on the European mainland.

Details on trapping and tagging, processing of raw monitoring data, date of last detection in the UK and arrival on the mainland of each individual can be found in supplementary file 1.

### 2.2 Acoustic monitoring

We performed continuous acoustic monitoring at 13 offshore platforms in the southern North Sea (figure 1), using an UltraSoundGate 116Hnbm system with an electret ultrasound microphone FG-DT50 (Avisoft Bioacoustics, Germany), installed at an average height of 21 m above sea level. Monitoring was conducted from March until June during four consecutive years (2018-2021). The effective monitoring effort included 4309 nights in total, respectively 1037, 1517, 914 and 841 per year. After retrieving and processing the raw monitoring data we removed the other bat species from the dataset (2.7% of the recordings), using R version 4.5.1 (R Core Team, 2025) and R studio (R-Studio Team, 2025). The dataset included 2777 Nathusius’ pipistrelle recordings of which 152 (5.5%) contained feeding buzzes/intense exploration calls (cf Barataud, 2020). To visualize the monitoring results, we made date-time plots using the R-package ggplot2 (Wickham, 2016) in combination with gghourglass (de Vries, 2025), in which the recorded bat activity is shown throughout the season and the night. Details on software settings, installation of the equipment, monitoring locations, monitoring periods and the processing of raw monitoring data can be found in supplementary file 2.

In the acoustic dataset, we assumed that the recorded bat activity at a particular moment refers to one individual bat as multiple bats simultaneously present at an offshore location in the southern North Sea has been documented only once (Lagerveld et al., 2014), and is therefore considered rare. Using an interval of two hours without bat activity to separate individuals resulted in 172 distinguished individual bats. We checked the sensitivity of this assumption by using alternative timeframes of 30 min, 60 min, four hours and one night. This resulted in an increase of 17 and 11 bats, and a decrease of 5 and 17 bats respectively. Thus, applying alternative timeframes resulted in a slight difference in the distinguished number of individuals (max 10 %). For each individual, we determined the first and last recorded occurrence and calculated the total duration it was detected at the offshore platform.

### 2.3 Weather data and lunar phase

All weather data were downloaded (2 July 2025) from the Climate Data Store of Copernicus (Hersbach et al., 2018), using the appropriate application user interface (https://cds.climate.copernicus.eu/how-to-api). Data were downloaded as a spatial raster (0.25×0.25°) with an hourly interval. For the analyses, data were selected from the raster cell located at 52.35° N 3.15° E (figure 1), which lies in the southern North Sea approximately halfway between Southwold in the UK (52.33° N 1.68° E) and Zandvoort in the Netherlands (52.37° N 4.53° E).

Several weather parameters were used in the analysis, including temperature at 2 m [K], total precipitation [m/h], eastward (U) and northward (V) wind speed components [m/s]. Wind speeds (U- and V-components) were provided for specific air pressure layers, which were converted to altitude layers using the barometric equation (NASA, 1976). To match the relatively large geographical scale of this study we used wind data at 200 m above sea level, as winds above the surface layer (up to 100-150 m) more closely represent regional airflow patterns (Garratt, 1992).

We calculated the atmospheric pressure change by subtracting the atmospheric pressure on a particular day with the atmospheric pressure from the previous day, and derived the temperature in degrees Celsius by subtracting 273,15 from the temperature in Kelvin. To reflect weather conditions at departure we took the mean value of each parameter in the time-intervals between sunset and two hours after sunset. The lunar phase [radians] was obtained using the R-package *suncalc* (Thieurmel, 2022).

### 2.4 Analysis

We compared seasonal and nightly bat occurrence of the tracked bats and the acoustically detected individuals. Subsequently, we analyzed departure decisions in relation to environmental conditions. For the second analysis we modelled the probability of bat migratory departures for both acoustic and tracking data within a similar analytical framework. ChatGPT was used to assist in drafting and debugging parts of the statistical analysis code (R/JAGS). No AI-based tools were used to generate results or interpret outputs. All code and outputs were critically reviewed, tested, and validated by the authors.

#### 2.4.1 Seasonal and nocturnal bat occurrence

The single night crossings from the tracking dataset with known departure time from the east coast of the UK and the arrival time at the Dutch coast (n=8) were used to examine the departure times relative to sunset, arrival times relative to sunrise, and the overall duration of the crossing. We compared the timing of occurrence in the night of the tracked bats with the acoustically recorded bats. In addition, we investigated whether acoustic bat presence in the night depends on seasonality and the spatial distribution. For this, we applied a generalized additive model (GAM) in the R-package mgcv (Wood, 2017), with time after sunset (when the bat was recorded for the first time) as the response variable. Night in year was entered as a low-rank thin-plate smoother to capture the seasonal-temporal pattern and latitude and longitude were included in a tensor product smoother to assess spatial differences. We made residual diagnostic plots and variograms to check for violations of the model assumptions.

#### 2.4.2 Environmental predictors

From the tracking dataset, we selected the individuals detected on the European mainland, thereby confirming migration from the UK over the North Sea (n=27). The night of the crossing over the North Sea was known for 10 individuals (of which the actual departure and arrival time were recorded for eight bats). The departure night was unknown for the remaining 17 individuals. To account for uncertainty in migration dates, we used a state-space modelling approach (Auger-Méthé et al., 2021; Baldwin et al., 2018). We estimated the probability of migratory departure across nights in a capture-recapture model in state-space formulation (Gimenez et al., 2007; Royle, 2008), structurally analogous to the nest survival models described in (Kéry and Schaub, 2011). In our model, migratory state was introduced as a matrix of individual ‘encounter histories’ throughout the migration period. The model estimated the probability that an individual transitions from a “non-migratory” state on night *t-1* to a “migratory” state on night *t*. This transition probability was modelled as a logistic function of environmental predictors. The ‘encounter history’ of each individual started with the date, when the individual was tagged. For each night, the state was classified as 1 (not yet departed, i.e. between the moment of tagging and the last detection in the UK), 0 (known to have departed from the UK), or NA (state is unknown, particularly, on the nights between the last registration in the UK and the first registration in continental Europe). This model setup allowed estimation of latent probability of migration (otherwise, transition from state 1 to state 0) for the nights with unobserved state.

To treat the acoustic dataset in the same modeling setup as the tracking data, we created encounter histories for each individual bat. During data exploration, we observed that acoustically recorded bats in May and June often coincided with westerly winds, cf. the pattern found in bats tracked from the UK (Lagerveld et al., 2024), while in March-April detections were mainly associated with southerly winds. We therefore assigned the individual bats to either the March-April or May-June period, based on the date of detection. For bats from the March-April period, encounter histories started from the night before the first ever bat was recorded at sea (15^th^ of March), or one day before the acoustic detector became operational later in the season. For individuals from the May-June period, encounter histories started on the 29^th^of April or one day before the detector was turned on. A total of 20 individual bats were excluded from the analysis because their pattern of occurrence across consecutive day and night periods indicated that they were potentially the same individuals detected before and after their diurnal stopover (cf Lagerveld et al., 2023b). These bats were detected around sunset (from one hour before to two hours after sunset) following detections during the previous night within three hours of sunrise. The estimated departure of the remaining bats recorded around sunset was allocated to the previous night, all other bats were allocated to the night in which they were recorded.

The initial fully-parameterized acoustic model included night number (linear and quadratic), windspeed (linear and quadratic), harmonic terms representing wind direction (cos θ, sin θ, cos 2θ, sin 2θ). Assuming that preferred windspeeds and migration periods (March-April or May-June) may differ, depending on the wind direction, we included interactions between each harmonic term and windspeed as well as period. Because temperature was correlated with night number, we used the residuals of temperature (on date) as linear covariate in the model. Atmospheric pressure change and cloud cover were also included as linear covariates, while precipitation was modeled as a binary covariate. The effect of lunar phase was represented by two sets of harmonic covariates, sine and cosine of the lunar phase (sinθ, cosθ) and their second-order harmonics (sin2θ, cos2θ).

The initial fully-parameterized tracking model included night number (linear and quadratic), windspeed (linear and quadratic), harmonic terms representing wind direction (cos θ, sin θ, cos 2θ, sin 2θ) and interactions between each harmonic term and windspeed. We did not test additional covariates in the tracking model to avoid overparameterization. In both models individual-level random intercepts were included to account for repeated observations within the encounter histories of the same bats. The standard deviation of the individual random effects was assigned a weakly informative half-Cauchy prior with scale 3.

We simplified the models by sequentially removing covariates with the weakest support, based on whether the 95% posterior credible interval of their beta parameters overlapped zero and on the corresponding evidence ratio (ER). Convergence was confirmed for all parameters (Rhat < 1.01). The effects of the statistically significant covariates were visualized using R-package ggplot2 (Wickham, 2016).

## 3 RESULTS

### 3.1 Seasonal and nocturnal occurrence

The night of the crossing was known for 10 tracked bats, of which the actual departure time from the UK coast and subsequent arrival time on the Dutch coast was recorded for eight individuals. These bats departed about one hour after sunset and arrived approximately three hours before sunrise at the European mainland. The sea crossing took three to five hours to cover a median distance of 172 km at a median minimal groundspeed of 43 km/h (table 1 and figure 2).

**Table 1:**
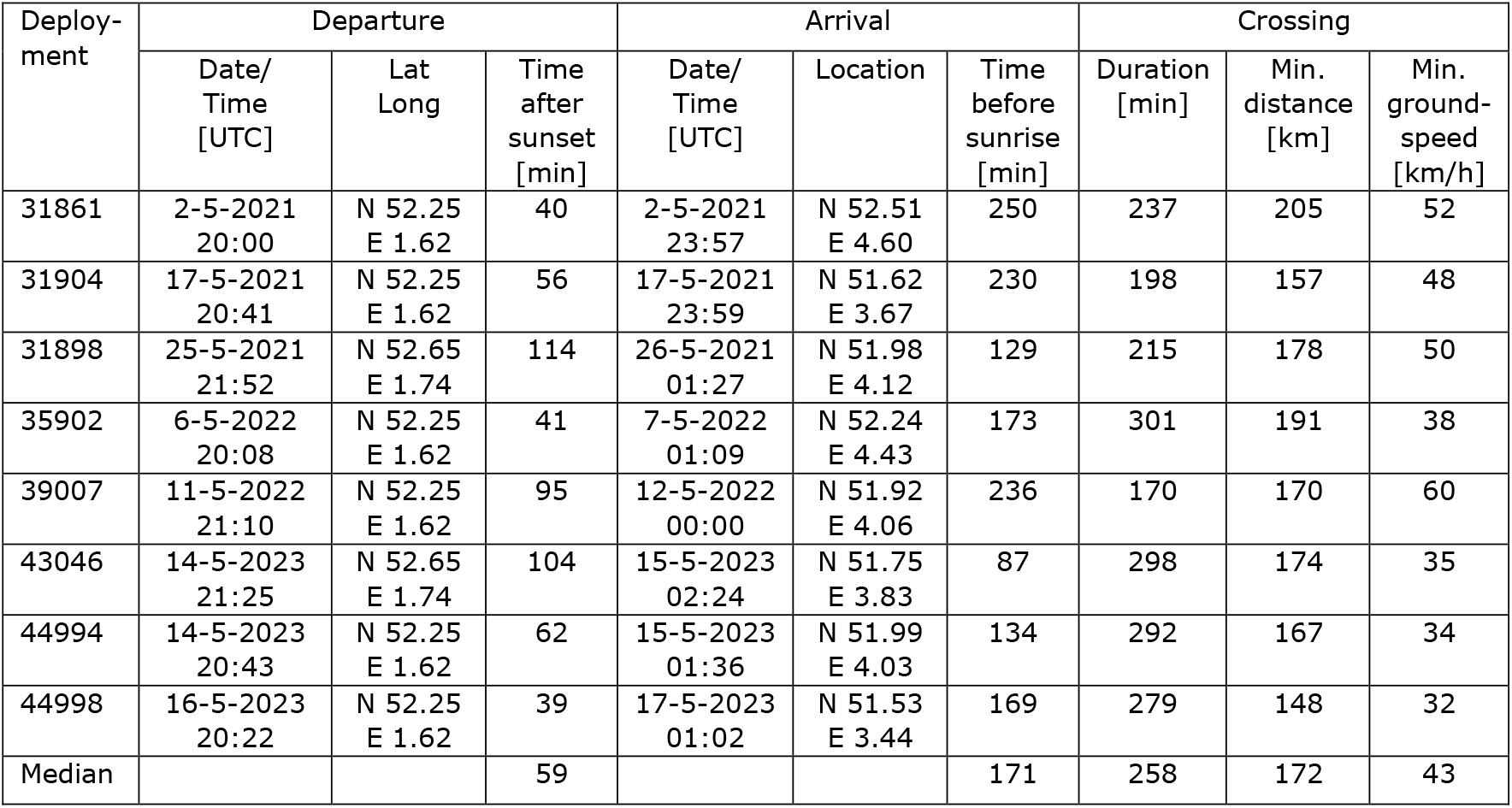
Departures and arrivals of single night crossings over the southern North Sea.

**Figure 2.**
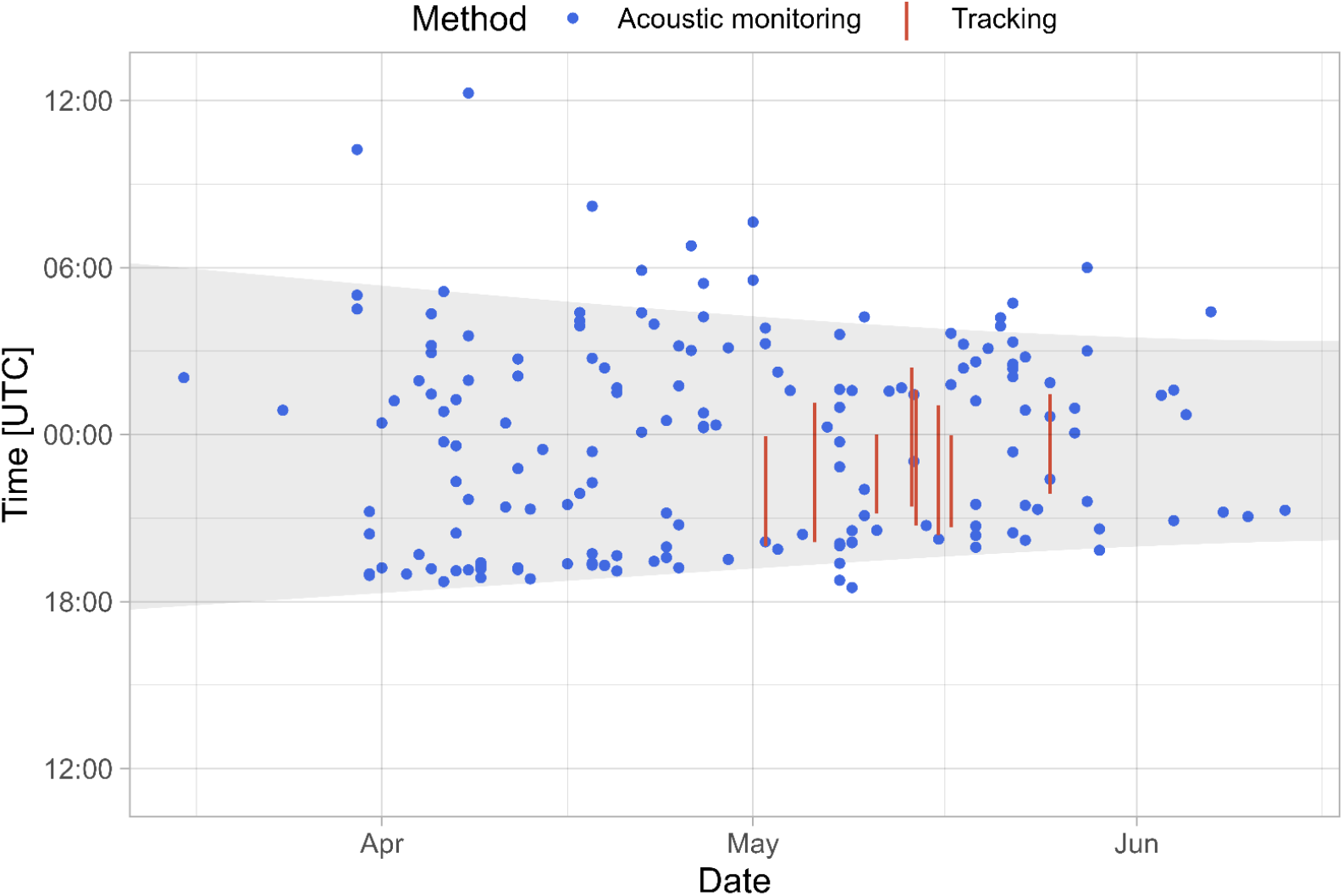
Acoustically recorded bat occurrence throughout the season for all monitoring locations and monitoring years (2018-2021). The time interval between sunset and sunrise is represented by grey. The dots represent the first recording of each individual (n=172). The red bars indicate the duration of single-night crossings over sea from the tracked individuals with known departure time from the east coast of the UK and arrival time at the coast of the European mainland (n=8). Note that the acoustically recorded bats refer to observations in Dutch and Belgian waters, while the crossings over sea include the timeframe of travel from the UK to the European mainland.

The earliest migrant amongst the acoustically recorded bats was observed 15 March 2020 (night number 75) and the last 12 June 2021 (night number 163). Migration peaked in April (night number 91-120) and May (night number 121-150), with 50% and 39% of the individuals (n=172) respectively. Bats were recorded at all monitoring locations, though not every year. Figure 2 shows the overall bat occurrence throughout the season and the night.

Bats were generally acoustically recorded in the night and sometimes during daylight hours. Most bats were recorded around sunset and sunrise: 37% within one hour before until two hours after sunset, 31% within three hours before sunrise and 9% were recorded during daylight hours, up to 7 hours after sunrise. The remaining 23% were recorded between two hours after sunset and three hours before sunrise (a time interval of three to seven hours, depending on the night number). When bats were recorded within one hour before until two hours after sunset (n=64), in 28% of the cases (n=18) bats were recorded the previous night three hours before sunrise (n=11, out of 46), or after sunrise (n=7, out of 15). The arrival time after sunset did not depend on the night number (p=0.450), nor the spatial location (p=0.173).

Most individuals (79%) were recorded less than 15 min: 103 bats (60%) less than 1 min, 16 bats (9%) between 1 – 5 min and 17 bats (10%) between 5-15 min. Longer observed individuals included 23 bats (13%) between 15-60 min, 6 bats (4%) between 1-2 hour, 3 bats (2%) between 2-4 hour and 4 (2%) bats were recorded more than 4 hours.

Detailed information on the yearly acoustic bat presence, occurrence throughout the season and the night, as well as the annual recorded bat activity for each monitoring location can be found in supplementary file 3.

### 3.2 Environmental predictors

In the simplified tracking (Eq. 1) and acoustic (Eq. 2) model, the effects of night number, linear and quadratic windspeed were retained, suggesting an increase in departure probability over the migration period and under moderate windspeeds. The effects of wind directions (expressed as harmonic terms) and speeds on the probability of departures differed between tracked bats and the two periods of acoustically detected bats.

The simplified tracking model was:

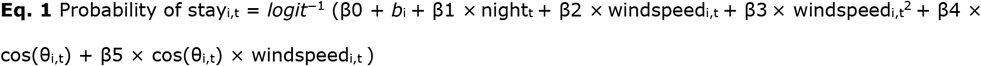

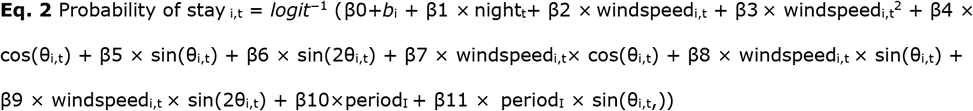

where

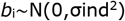

are the individual-specific random effects.

It should be noted that, in the model output (supplementary file 4), the probability of stay is modelled. A negative coefficient in the linear predictor therefore increases the probability of departure.

In Figure 3, we summarize wind speeds and the relative predicted migration probabilities for observed departures in the tracking model and in each period of the acoustic model. The baseline migration probability differed between the periods, being higher in the May–June period due to the shorter encounter histories. In the figure, we present therefore relative migration probabilities within each period. Specifically, for each directional bin, the median predicted probability is scaled by the sum of median probabilities across all bins, allowing comparison of wind effects independently of baseline differences.

**Figure 3.**
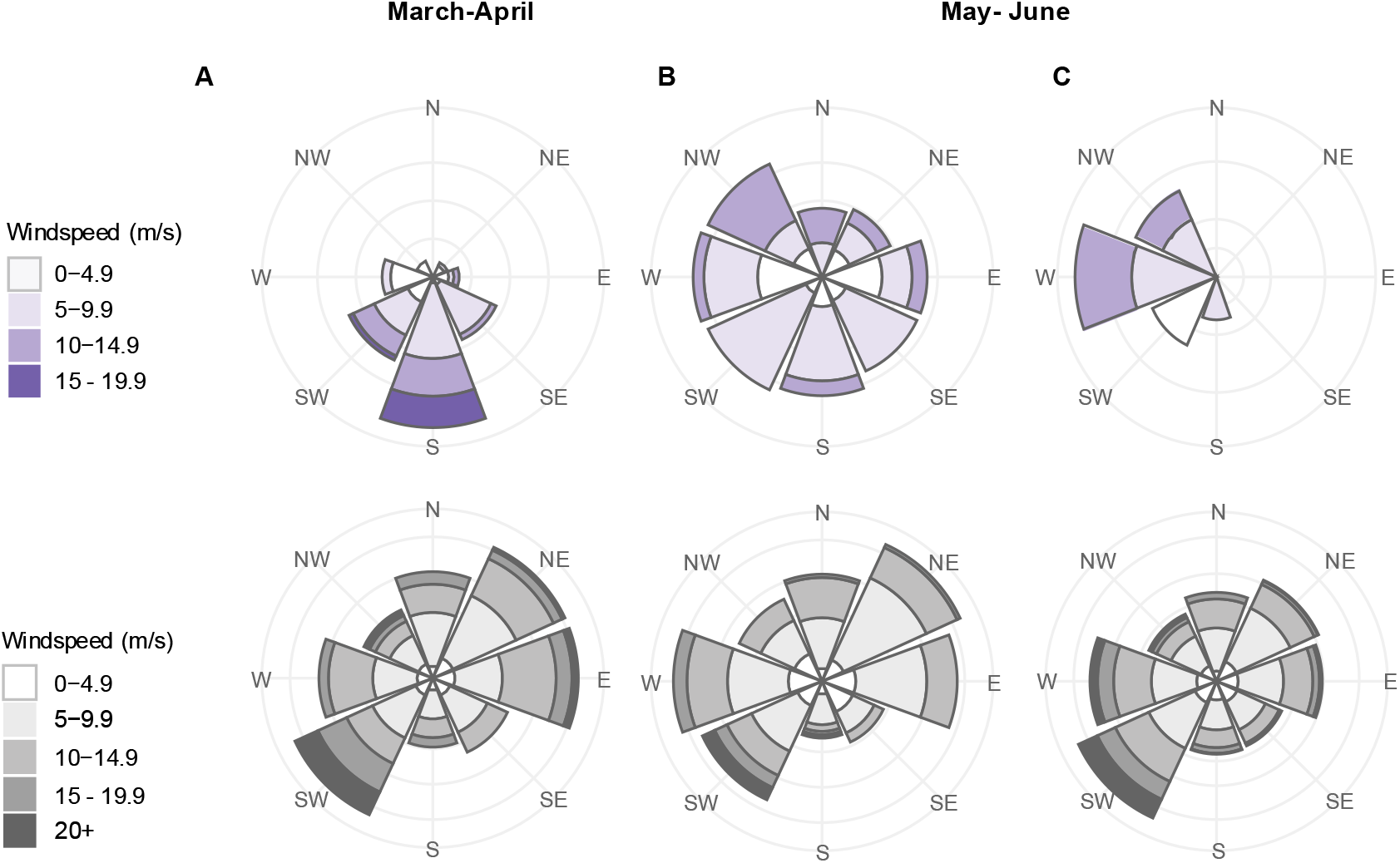
Predicted probability of spring migration over the North Sea as a function of wind conditions at 200 meter above sea level (upper row) and prevailing wind conditions throughout the migration period (lower row) for (A) the acoustic dataset in March–April and (B) May–June, and (C) for the tracking dataset, where migration took place in May–June. The upper-row wind roses represent the directional structure of relative migration probability for each period. Meteorological wind direction (direction from which the wind blows) is divided into 8 compass sectors (45° bins). For each sector, the total bar height represents the median predicted probability of departure, normalized within period. Bars are stacked by wind-speed class, with the purple fill intensity indicating the proportion of observations falling into each wind-speed class within a given direction. The wind roses in the lower row illustrate the directional structure of overall winds during the corresponding migration periods. The height of each bar indicates the proportion of nights with winds originating from each directional bin. Stacks within each bar indicate the proportional distribution of wind-speed classes within each direction. Shades of grey represent wind-speed classes, with lighter shades denoting weaker winds.

Departures of tracked bats occurred throughout May and June. The model predicted high co-occurrence of departures with moderate westerly winds. Of the 10 individuals with known departure night 8 were observed departing with winds from west or northwest (median windspeed = 7.0 m/s, quartile range: 5,4 to 11.2 m/s; Fig. 3).

In the acoustic model (Eq. 2), the role of westerly winds was also prominent in the departures recorded in May-June. For bats recorded under winds from west and northwest, median windspeed was 6,6 m/s, quartile range: 3,0 to 9.9 m/s; n=32; Fig. 3). Bats were also observed in southerly and easterly winds. For bats recorded under winds from southwest, south and southeast, median windspeed was 5.9 m/s; quartile range: 5.2 to 7.5 m/s; n=19; Fig. 3) and from east, median windspeed was 3.4 m/s; quartile range 3.4-5.6 m/s; n=7)

Bats from the March-April period were most likely to be detected over the North Sea in stronger winds from southerly directions (median windspeed = 8.5 m/s, quartile range: 5.9 to 12.7 m/s for the observed departures in winds from southwest, south and southeast; n=68). For both periods, predicted migration was unlikely under winds from north and northeast (Fig. 3).

All other environmental covariates (lunar phase, cloud cover, precipitation, atmospheric pressure change and temperature) showed no statistically significant effect. The output of the analysis can be found in supplementary file 4.

## 4 DISCUSSION

In the present study we combined two observation techniques, acoustic monitoring and individual tracking, to improve our understanding of bat migration patterns. Spring tracking data indicated that bats departed the UK’s east coast approximately one hour after sunset and reached the Dutch coast about three hours before sunrise, whereas acoustic monitoring showed that most offshore detections occurred close to sunset or sunrise. Both monitoring methods identified wind conditions at 200 m above sea level as a key predictor of Nathusius’ pipistrelle spring migration over sea. Tracked bats crossed from the UK to the European mainland in May and June under supportive moderate westerly and north-westerly winds. Similarly, most acoustically recorded bats in May–June were also detected under westerly and north-westerly winds, albeit at lower wind speeds. However, acoustically recorded bats were absent in March–April under these wind conditions. During this earlier period, most acoustic detections were associated with strong southerly winds, while no tracked bats were observed crossing the North Sea. No significant effects were detected for the other environmental variables examined, including precipitation, temperature, cloud cover, atmospheric pressure change and lunar phase.

### 4.1 Seasonal and nocturnal occurrence

The main breeding areas of Nathusius’ pipistrelle are located in northeastern Europe and wintering occurs in southern and western Europe, including the UK (Russ, 2022). Migrants generally follow a southwest– northeast route in spring, reversing direction in autumn (Hutterer, 2006; Pētersons, 2004). North Sea crossings from the east coast of the UK to the European mainland are typically routed east to southeast to reduce the over-water distance (Lagerveld et al., 2024).

Our data show that spring migration over the southern North Sea occurs from mid-March until mid-June, and most bats can be expected in April and May (median acoustic occurrence 27 April). The species has been recorded in the German Bight between 20 April and 26 May (median 2 May) (Hüppop and Hill, 2016) and migration over central and northern Europe occurs mainly in May, with the median observation early May in northwest Germany (N 54°) and late May in southwest Finland (N 60–61°) with a difference of 20 days between the two areas (Rydell et al., 2014). Therefore, the seasonal pattern we captured fits well in the pattern from other studies.

The timing of bat occurrence throughout the night differed significantly between the tracking and acoustic datasets. Tracked bats that crossed the North Sea in a single night departed from the UK’s east coast about one hour after sunset and arrived at the Dutch coast approximately three hours before sunrise. The sea crossing took around four hours. In contrast, acoustically recorded bats were mostly detected around sunset or sunrise. Since there are no resident bat populations at the North Sea, detections near sunset indicate that bats used offshore platforms as diurnal stopover sites during their migration over sea (Lagerveld et al., 2023b). Late-night detections, or morning occurrences in daylight hours likely represent individuals seeking daytime shelter.

Another indication that offshore platforms are primarily used as roosting sites, and not as foraging sites, is the generally short duration bats are recorded acoustically and the low proportion of recordings containing feeding buzzes/intense exploration calls.

### 4.2 Environmental predictors

Supportive winds promote bat migratory departures (Dechmann et al., 2017; Hurme et al., 2025; Lagerveld et al., 2024, 2021). Consistent with these studies, our analysis identifies wind at 200 m above sea level as a key determinant of bat migratory movements. Moreover, seasonal differences in the wind conditions under which bats were recorded suggest migration from distinct geographical areas. In March– April, acoustic detections coincided with relatively strong southerly winds (median wind speed = 8.5 m/s, interquartile range: 5.9–12.7 m/s), generally exceeding the maximum range velocity of Nathusius’ pipistrelle (7.5 m/s, cf Troxell et al., 2019), which represents the species’ typical migratory airspeed. In these conditions a migrating animal has no other option than to follow the general direction of the wind (Alerstam, 1978). Consequently, early spring detections most likely reflect bats that were wind-drifted over the North Sea while migrating from areas south of the acoustic monitoring network (e.g. Channel Area, northern France, Belgium, and southern parts of the Netherlands). Migration from the east coast of the UK to the European mainland in March–April appears to be rare or absent, as none of the tagged bats migrated over sea in this period and acoustic detections during westerly and northwesterly winds were scarce.

All tagged bats crossed the North Sea in May–June, typically under moderate westerly or northwesterly tailwinds (median wind speed = 7.0 m/s, interquartile range: 5.4–11.2 m/s). Similarly, acoustic detections in May–June were predominantly associated with westerly and northwesterly winds, albeit at lower wind speeds (median = 6.6 m/s, interquartile range: 3.0–9.9 m/s), indicating reduced wind support for bats detected acoustically. During May-June, migration probability was also elevated under winds from southerly directions (median = 5.9 m/s; interquartile range: 5.2–7.5 m/s) and from the east (median = 3.4 m/s; interquartile range: 3.4–5.6 m/s). Under these conditions, wind-drifted bats originating from the European mainland appear less likely, instead, it seems more plausible that these observations consist of individuals departing from the UK under crosswinds or low headwinds. In both periods, northerly and northeasterly winds were associated with the lowest migration probabilities.

In addition to wind, several other environmental factors such as precipitation, cloud cover, atmospheric pressure change, temperature and the lunar phase, can influence bat migratory movements. Increased temperatures (Hurme et al., 2025; Pettit and O’Keefe, 2017; True et al., 2021) and decreasing atmospheric pressure (Hurme et al., 2025) trigger spring migration. During autumn migratory departures generally occur under dry conditions (Brabant et al., 2021; Lagerveld et al., 2014, 2023b), whereas precipitation and overcast skies tend to induce stopovers (Cryan and Brown, 2007; Hüppop and Hill, 2016). Autumn migration has also been linked to darker lunar phases (Cryan and Brown, 2007; Lagerveld et al., 2023b). Our analysis of the acoustic dataset revealed no statistically significant effect of precipitation, cloud cover, atmospheric pressure change, temperature and lunar phase. One possible explanation is that bats acoustically recorded on offshore platforms may have landed under a wide range of environmental conditions, including those that are often considered suboptimal for migration.

### 4.3 Methodological implications

Our novel analytical approach – a state-space formulation of the capture-recapture model – has proven to be an effective solution for analyzing bat departure decisions. It enabled the comparable analysis of two datasets collected using different monitoring techniques, and incorporated uncertainties in migration dates of tagged individuals. While these observation techniques were thought to provide contradictory insights into migration patterns, our analysis shows that combining these two observation techniques improves our understanding of bat migration patterns.

Although the acoustic and tracking monitoring results capture the same late spring seasonal migration pattern, both methods have certain limitations and therefore provide information on different fractions of migratory bats. Acoustic detectors at offshore platforms record Nathusius’ pipistrelles below elevations of 50 m above sea level (Lagerveld et al., 2023b) and seem to primarily record individuals that landed on these structures as they failed to cross the North Sea in one attempt, e.g. due to less favorable wind conditions (reduced tailwinds, crosswind or headwind), or because they departed from locations further inland. In contrast, tracking data largely represented bats that crossed the North Sea under moderate supportive tailwinds in a single flight. During supportive tailwinds bats generally fly at altitudes too high to be detected acoustically (Lagerveld et al., 2024). Consequently, bats benefiting from stronger tailwinds, may be underrepresented -or even absent-in the acoustic dataset. This implies that curtailment measures relying exclusively on acoustic monitoring (cf Rijkswaterstaat, 2021) underestimate bat migration over sea at higher wind speeds. Tracking data also have limitations: they reflect only bats departing from a specific geographic location -in this study the east coast of the UK - and therefore remain blind to the migration originating from other areas. To reveal the geographic origin of the Nathusius’ pipistrelles appearing over the North Sea with southerly winds will require extensive tracking studies of individuals captured in other regions such as the Channel area, Northern France and Belgium.

### 4.4 Mitigation strategies for offshore wind farms

Although detection ranges of acoustic detectors and Motus receivers differ, both the acoustic and tracking models indicated that wind at 200 m above sea level (representing regional airflow) was a strong determinant of migration probability. Since current models for curtailment of offshore wind farms require environmental parameters that best predict migration (Rijkswaterstaat, 2021), we can recommend wind direction and speed at 200 m above sea level as key candidate parameters for tailoring curtailment measures to protect Nathusius’ pipistrelle during spring migration over the southern North Sea.

Moreover, acoustic monitoring can capture migration originating from different geographic locations but likely underestimates the part of the population that crosses the North Sea with optimal wind support in a single flight. This suggests that acoustic and tracking data are complementary rather than contradictory, and both monitoring methods should be used together when developing mitigation measures for offshore wind farms.

## AUTHOR CONTRIBUTIONS

Sander Lagerveld conceived the ideas and designed the study. Jane Harris, Bart Noort and Sander Lagerveld conducted the fieldwork and collected the data. Julia Karagicheva, Sander Lagerveld, Pepijn de Vries and Eldar Rakhimberdiev analysed the data. Sander Lagerveld and Julia Karagicheva led the writing of the manuscript. Sander Lagerveld, Jane Harris, Martin Poot and Frank van Langevelde acquired funding. All authors contributed critically to the drafts and gave final approval for publication.

## ETHICS DECLARATIONS

Bat trapping and tagging was carried out under Natural England project licences 2021-55582-SCI and 2022-63237-SCI

## DATA AVAILABILITY STATEMENT

Data and analysis scripts are available from Zenodo: https://doi.org/10.5281/zenodo.18485034 (Lagerveld & Karagicheva 2026)

## ACKNOWLEDGEMENTS

We sincerely thank Nathan Duszynski, Lotty Packman, Ewan Parsons, Sue Parsons and Huma Pearce for their invaluable help in the field during bat tagging activities. The installation and maintenance of the Motus receiver network was carried out primarily by Martijn Keur and Cor Sonneveld in the Netherlands, René Janssen in Belgium, Ewan Parsons in the UK and Heinz-Hinrich Blikslager, Thomas Klinner, Thomas Mertens, Mario de Neidels, and Florian Packmor in Germany. We also like to thank Meik Verdonk and Henri Zomer (Dutch Ministry of Infrastructure and Water Management) for their valuable comments and suggestions, which greatly improved this manuscript.

