## Supplementary file 1 for "Directional variation and method-specific detection patterns in offshore bat migration: implications for wind farm mitigation"

**Supplementary file 1: Trapping and tagging effort, processing MOTUS data and tracking results**

*Trapping and tagging*

Bats were trapped and tagged at Minsmere (N 52.24 E 1.62), Benacre National Nature Reserve (N 52.39 E 1.71) and the Landguard Bird Observatory (N 51.94 E 1.32). In total 36 trapping sessions were undertaken, lasting on average 4.5 hours (Table SF1-1).

*Table SF1-1: trapping sessions*

| Date | Location | Start time [UTC] | End time [UTC] | Duration [h] | Harp traps | Mist nets | Team* |
|---|---|---|---|---|---|---|---|
| 29-03-21 | Minsmere | 18:25 | 22:00 | 03:35 | 2 | 3 | JH, SP, EP |
| 30-03-21 | Minsmere | 18:35 | 22:30 | 03:55 | 2 | 2 | JH, SP, EP |
| 04-04-21 | Minsmere | 18:45 | 21:30 | 02:45 | 2 | 2 | JH, SP, EP |
| 18-04-21 | Minsmere | 19:30 | 21:30 | 02:00 | 2 | 3 | JH, SP, EP |
| 19-04-21 | Minsmere | 19:20 | 21:00 | 01:40 | 2 | 2 | JH, SP, EP |
| 20-04-21 | Minsmere | 19:15 | 23:00 | 03:45 | 2 | 3 | JH, SP, EP |
| 27-04-21 | Minsmere | 19:15 | 23:34 | 04:19 | 3 | | JH, SP, EP |
| 02-05-21 | Minsmere | 19:30 | 23:00 | 03:30 | 3 | | JH, SP, EP |
| 07-05-21 | Minsmere | 19:30 | 21:55 | 02:25 | 3 | | JH, SP, EP |
| 08-05-21 | Minsmere | 19:30 | 02:00 | 06:30 | 3 | | JH, SP, EP |
| 10-05-21 | Minsmere | 19:35 | 00:30 | 05:00 | 3 | | JH, SP, EP |
| 11-05-21 | Minsmere | 19:35 | 02:00 | 06:30 | 3 | | JH, SP, EP |
| 12-05-21 | Minsmere | 19:40 | 23:45 | 04:05 | 3 | | JH, SP, EP |
| 14-05-21 | Minsmere | 19:40 | 02:00 | 06:20 | 3 | | JH, SP, EP |
| 05-04-22 | Minsmere | 19:40 | 23:00 | 03:20 | 3 | | JH, SP, EP |
| 13-04-22 | Minsmere | 19:00 | 23:50 | 04:50 | 3 | | JH, SP, EP |
| 14-04-22 | Minsmere | 19:00 | 00:30 | 05:30 | 3 | | JH, SP, EP |
| 18-04-22 | Minsmere | 19:00 | 00:33 | 05:33 | 3 | | JH, SP, EP |
| 19-04-22 | Minsmere | 19:15 | 22:00 | 02:45 | 3 | | JH, SP, EP, HP |
| 30-04-22 | Minsmere | 19:30 | 00:00 | 04:30 | 5 | | JH, SP, EP, HP, ND |
| 01-05-22 | Minsmere | 19:30 | 23:30 | 04:00 | 5 | | JH, SP, EP, HP, ND |
| 04-05-22 | Landguard | 19:35 | 00:30 | 04:25 | 3 | | JH, SP, EP, HP |
| 05-05-22 | Landguard | 19:35 | 00:30 | 04:25 | 3 | | JH, SP, EP, HP |
| 10-05-22 | Minsmere | 19:50 | 01:30 | 05:40 | 3 | | JH, SP, EP, HP |
| 30-03-23 | Minsmere | 18:25 | 21:55 | 03:30 | 4 | | JH, HP, ND |
| 17-04-23 | Minsmere | 19:00 | 21:00 | 02:00 | 4 | | JH, HP, ND |
| 22-04-23 | Minsmere | 19:05 | 23:30 | 04:25 | 5 | | JH, HP, ND |
| 28-04-23 | Minsmere | 19:20 | 01:35 | 06:15 | 5 | | JH, HP, ND |
| 29-04-23 | Benacre | 19:20 | 00:30 | 05:20 | 5 | | JH, HP, LP |
| 30-04-23 | Minsmere | 19:30 | 02:30 | 07:00 | 5 | | JH, SP, EP, HP |
| 01-05-23 | Minsmere | 19:30 | 02:00 | 06:30 | 5 | | JH, SP, EP, HP |
| 02-05-23 | Benacre | 19:30 | 22:30 | 03:00 | 5 | | JH, SP, EP, HP |
| 03-05-23 | Minsmere | 19:30 | 02:30 | 07:00 | 5 | | JH, SP, EP, HP |
| 05-05-23 | Benacre | 19:30 | 01:00 | 05:30 | 5 | | JH, SP, EP, HP |
| 07-05-23 | Minsmere | 19:30 | 02:30 | 07:00 | 5 | | JH, SP, EP, HP, ND |
| 10-05-23 | Minsmere | 20:00 | 02:30 | 06:30 | 5 | | JH, SP, EP, HP |

* JH = Jane Harris, SP = Sue Parsons, EP = Ewan Parsons, HP = Huma Pearce, ND = Nathan Duszynski, LP = Lotty Packman

Bats were captured with no. 2 and no. 3 Austbat harp traps (Faunatec, Australia) and mist nets (Solida, Germany) in combination with AT100 acoustic lures (Binary Acoustic Technology, United States) playing the advertisement calls of male Nathusius' pipistrelle. Traps were checked every 20 minutes, or sooner, mist nets were monitored continuously. Of each individual information on biometry (forearm length), sex, age (when possible) and reproductive status was gathered using the criteria described by Haarsma, 2008. Subsequently, a visual check for pregnancy, injuries, parasites or aberrations in fur and wing membranes was executed. Pregnant females, judged to be less than 2 weeks to parturition, were released immediately. Individuals lacking visible injuries, low ectoparasite load, healthy fur and wing membranes were qualified suitable for tagging. They were tagged only, when their weight exceeded 6 g. This minimum was employed, in accordance with Natural England guidance for the capture and marking of bats (guidance note WML-G39), to ensure that the weight of the radio-transmitter should not exceed 5% of the individual's body weight. To attach the radio-transmitter, fur was lightly clipped between the shoulder blades to create a small pocket (8 mm x 4 mm). The transmitter was affixed using medical adhesive (Torbot Group, United States). To ensure the transmitter was properly attached, tagged bats were held in cloth bags on heat pads and released as soon as the adhesive was set (after about 10 min). A total of 139 Nathusius' pipistrelles were trapped, of which 122 individuals were tagged.

*Processing of MOTUS data*

Receiver metadata and transmitter detection data were retrieved (19 May 2025) from www.motus.org, using the Motus R package (Birds Canada, 2022). Detections with run lengths < 3 were filtered out, in accordance with the MOTUS R Book (Birds Canada, 2022). In addition, unlikely detections were removed from specific receivers at locations with high levels of radio noise (table SF1-2). The location of a receiver that detects the radio transmitter is used as an estimate for the bat's current location. If multiple receivers detect a signal from a particular transmitter within 30 seconds and are less than 10 km apart, the receiver with the strongest signal is considered the best approximation of the bat's location. When a bat moved from one receiver to another we assigned the departure and arrival timestamps when the strongest signals were received.

*Tabe SF1-2: false positives*

| Deployment | Detections removed | Reason |
|---|---|---|
| 43047 | all detections<br>17_Fedderwardersiel (#11191) | A receiver in an area with a lot of radio activity nearby. Lots of detections during a prolonged time, also during daylight hours and from other deployments. |
| 43048 | 18/5/2023<br>26_Utlandshörn (#10968) | Detections during daylight hours |
| 43052 | all detections<br>17_Fedderwardersiel (#11191) | A receiver in an area with a lot of radio activity nearby. Lots of detections during a prolonged time, also during daylight hours and from other deployments. |
| 43046 | all detections<br>17_Fedderwardersiel (#11191) | A receiver in an area with a lot of radio activity nearby. Lots of detections during a prolonged time, also during daylight hours and from other deployments. |
| 42442 | all detections<br>17_Fedderwardersiel (#11191) | A receiver in an area with a lot of radio activity nearby. Lots of detections during a prolonged time, also during daylight hours and from other deployments. |
| 44997 | all detections<br>123_Norderney 2 (#10283) | Lots of detections during a prolonged time (also during daylight hours) |
| 31911 | all detections<br>WUR Texel NIOZ (#10258) | Multiple false positives of various species due the presence of many running (non-deployed) tags at the NIOZ office, very close to the receiver |
| 31912 | all detections<br>WUR Texel NIOZ (#10258) | Multiple false positives of various species due the presence of many running (non-deployed) tags at the NIOZ office, very close to the receiver |
| 31901 | all detections<br>Rysum 1 (#10282) | Multiple false positives of various species due the presence of many deployed tags in the immediate vicinity |
| 31907 | all detections<br>Rysum 1 (#10282) | Multiple false positives of various species due the presence of many deployed tags in the immediate vicinity |
| 31909 | all detections<br>125_Norderney 4 (#10285) | Multiple false positives of various species due the presence of many deployed tags in the immediate vicinity |
| 38491 | 12/6/2021<br>127_Woldsee 1 (#10555) | Multiple false positives of various species due the presence of many deployed tags in the immediate vicinity |
| 31902 | all detections<br>17_Fedderwardersiel (#11191) | A receiver in an area with a lot of radio activity nearby. Lots of detections during a prolonged time, also during daylight hours and from other deployments. |
| 39000 | all detections<br>17_Fedderwardersiel (#11191) | A receiver in an area with a lot of radio activity nearby. Lots of detections during a prolonged time, also during daylight hours and from other deployments. |
| 31905 | 19/5/2021 Minsmere2 (#12367)<br>21/5/2021<br>BM3-NPip_tracking (#11931)<br>23/5/2021 FB-Testing (#12097) | Tag found in Sparrowhawk pellet and retrieved. Remarkably it was still functioning (and not switched off) and subsequently detected by several receivers over the next few days during fieldwork activities. |

*Tracking results*

In total, 122 bats were tagged, comprising 18 males and 104 females. The number of individuals tagged differed across years, with 20 in 2021, 24 in 2022, and 78 in 2023. Table SF1-3 shows where and when each individual (deployment) was tagged, its last detection in the UK and the first documented arrival on the European mainland.

*Table SF 1-3: Time and location of tagging, last UK detection and first detection at the European mainland*

| Deployment | Sex | Tagged [UTC] | Lat/Long | Last UK [UTC] | Lat/Long | First mainland [UTC] | Lat/Long | Comments | Link |
|---|---|---|---|---|---|---|---|---|---|
| 31861 | F | 2021-03-29 21:50 | N 52.246 E 1.618 | 2021-05-02 20:13 | N 52.247 E 1.615 | 2021-05-02 23:57 | N 52.506 E 4.601 | crossing night 2021-05-02 | https://motus.org/data/tagDeployment?id=31861 |
| 31862 | M | 2021-04-20 21:00 | N 52.247 E 1.619 | 2021-06-04 00:55 | N 52.247 E 1.615 | | | | https://motus.org/data/tagDeployment?id=31862 |
| 31863 | M | 2021-05-02 21:55 | N 52.246 E 1.618 | 2021-06-04 23:14 | N 52.247 E 1.615 | | N 52.247 E 1.615 | | https://motus.org/data/tagDeployment?id=31863 |
| 31864 | M | 2021-05-08 20:55 | N 52.246 E 1.618 | 2021-06-13 22:52 | N 52.247 E 1.615 | | | | https://motus.org/data/tagDeployment?id=31864 |
| 31911 | F | 2021-05-08 22:25 | N 52.246 E 1.618 | 2021-05-12 20:50 | N 52.247 E 1.615 | | | | https://motus.org/data/tagDeployment?id=31911 |
| 31912 | F | 2021-05-08 23:55 | N 52.246 E 1.618 | 2021-05-12 23:25 | N 52.247 E 1.615 | | | | https://motus.org/data/tagDeployment?id=31912 |
| 31899 | F | 2021-05-09 00:20 | N 52.246 E 1.618 | 2021-05-10 21:17 | N 52.247 E 1.615 | | | | https://motus.org/data/tagDeployment?id=31899 |
| 31900 | F | 2021-05-10 22:05 | N 52.247 E 1.619 | NA | NA | | | tag found detached 2021-05-10 | https://motus.org/data/tagDeployment?id=31900 |
| 31901 | F | 2021-05-11 00:05 | N 52.247 E 1.619 | 2021-05-25 00:45 | N 52.247 E 1.615 | 2021-05-26 23:55 | N 51.116 E 26.324 | crossing night of 24, 25 or 26 May | https://motus.org/data/tagDeployment?id=31901 |
| 31902 | F | 2021-05-11 00:45 | N 52.247 E 1.619 | 2021-05-16 22:23 | N 52.247 E 1.615 | | | | https://motus.org/data/tagDeployment?id=31902 |
| 31903 | F | 2021-05-11 22:00 | N 52.247 E 1.619 | 2021-05-18 22:34 | N 52.247 E 1.615 | | | | https://motus.org/data/tagDeployment?id=31903 |
| 31904 | F | 2021-05-11 22:55 | N 52.247 E 1.619 | 2021-05-17 20:42 | N 52.247 E 1.615 | 2021-05-17 23:59 | N 51.623 E 1.619 | crossing night 2021-05-17 | https://motus.org/data/tagDeployment?id=31904 |
| 31905 | F | 2021-05-12 01:52 | N 52.247 E 1.619 | NA | NA | | | Predated | https://motus.org/data/tagDeployment?id=31905 |
| 31906 | F | 2021-05-12 22:15 | N 52.247 E 1.619 | 2021-05-29 00:41 | N 52.247 E 1.615 | | | | https://motus.org/data/tagDeployment?id=31906 |
| 31907 | F | 2021-05-13 00:35 | N 52.247 E 1.619 | 2021-05-19 21:28 | N 52.247 E 1.615 | | | | https://motus.org/data/tagDeployment?id=31907 |
| 31908 | F | 2021-05-14 22:45 | N 52.247 E 1.619 | 2021-05-25 21:20 | N 52.247 E 1.615 | | | | https://motus.org/data/tagDeployment?id=31908 |
| 31909 | F | 2021-05-15 00:15 | N 52.247 E 1.619 | 2021-05-22 23:28 | N 52.247 E 1.615 | | | | https://motus.org/data/tagDeployment?id=31909 |
| 31910 | F | 2021-05-15 00:35 | N 52.247 E 1.619 | 2021-05-18 21:25 | N 52.247 E 1.615 | 2021-05-20 21:22 | N 53.369 E 7.011 | first detection Germany early evening 2021-05-20. Consequently, crossing was in the night of 18 or 19 May | https://motus.org/data/tagDeployment?id=31910 |
| 31898 | F | 2021-05-15 01:40 | N 52.247 E 1.619 | 2021-05-25 22:01 | N 52.247 E 1.615 | 2021-05-26 01:22 | N 51.991 E 4.124 | crossing night 2021-05-25 | https://motus.org/data/tagDeployment?id=31898 |
| 31897 | F | 2021-05-15 20:30 | N 52.247 E 1.619 | 2021-05-24 21:55 | N 52.247 E 1.615 | 2021-05-28 21:07 | N 51.991 E 4.124 | first detection in Netherlands early evening 2021-05-28. Consequently, crossing was in the night of 24, 25, 26 or 27 May | https://motus.org/data/tagDeployment?id=31897 |
| 38458 | F | 2022-04-13 20:20 | N 52.247 E 1.619 | 2022-05-03 20:19 | N 52.247 E 1.615 | | | | https://motus.org/data/tagDeployment?id=38458 |
| 38459 | F | 2022-04-13 20:20 | N 52.247 E 1.619 | 2022-05-02 20:15 | N 52.247 E 1.615 | | | | https://motus.org/data/tagDeployment?id=38459 |
| 38460 | F | 2022-04-13 21:40 | N 52.247 E 1.619 | 2022-05-04 23:00 | N 52.247 E 1.615 | | | | https://motus.org/data/tagDeployment?id=38460 |
| 38491 | F | 2022-04-18 21:20 | N 52.247 E 1.619 | 2022-04-20 22:02 | N 51.940 E 1.327 | | | | https://motus.org/data/tagDeployment?id=38491 |
| 38492 | F | 2022-04-19 00:30 | N 52.247 E 1.619 | 2022-05-01 23:23 | N 52.247 E 1.615 | | | | https://motus.org/data/tagDeployment?id=38492 |
| 35904 | F | 2022-04-22 20:45 | N 52.247 E 1.619 | 2022-05-05 20:33 | N 52.400 E 1.727 | | | | https://motus.org/data/tagDeployment?id=35904 |
| 35902 | F | 2022-04-30 20:50 | N 52.247 E 1.619 | 2022-05-06 20:29 | N 52.247 E 1.615 | 2022-05-07 01:09 | N 52.247 E 4.433 | crossing night 2022-05-06 | https://motus.org/data/tagDeployment?id=35902 |
| 38862 | F | 2022-04-30 20:50 | N 52.247 E 1.619 | 2022-05-04 20:32 | N 52.247 E 1.615 | | | | https://motus.org/data/tagDeployment?id=38862 |
| 38863 | F | 2022-05-01 18:00 | N 52.247 E 1.619 | 2022-05-06 20:10 | N 52.247 E 1.615 | | | | https://motus.org/data/tagDeployment?id=38863 |
| 35903 | M | 2022-05-01 19:30 | N 52.247 E 1.619 | 2022-05-24 21:46 | N 52.247 E 1.615 | 2022-05-25 22:58 | N 51.632 E 2.765 | crossing night 2022-05-25 (detected at sea) | https://motus.org/data/tagDeployment?id=35903 |
| 38864 | F | 2022-05-01 20:45 | N 52.247 E 1.619 | 2022-05-01 23:31 | N 52.247 E 1.615 | | | | https://motus.org/data/tagDeployment?id=38864 |
| 38865 | F | 2022-05-01 21:30 | N 52.247 E 1.619 | 2022-05-06 23:26 | N 52.247 E 1.615 | | | | https://motus.org/data/tagDeployment?id=38865 |
| 38866 | F | 2022-05-01 22:00 | N 52.247 E 1.619 | 2022-05-02 00:19 | N 52.247 E 1.615 | 2022-05-06 21:02 | N 53.100 E 4.898 | crossing in the night of 2, 3, 4 or 5 May (crossing in the night of 1 May unlikely given the late detection , and early evening detection in the Netherlands on 6 May) | https://motus.org/data/tagDeployment?id=38866 |
| 38867 | F | 2022-05-01 22:00 | N 52.247 E 1.619 | 2022-05-03 01:48 | N 52.247 E 1.615 | | | | https://motus.org/data/tagDeployment?id=38867 |
| 38872 | F | 2022-05-01 22:25 | N 52.247 E 1.619 | 2022-05-06 20:52 | N 51.362 E 1.375 | 2022-05-09 22:43 | N 51.116 E 2.632 | crossing in the night of 6, 7, 8 or 9 May | https://motus.org/data/tagDeployment?id=38872 |
| 38868 | F | 2022-05-01 22:30 | N 52.247 E 1.619 | 2022-05-06 21:15 | N 52.247 E 1.615 | | | | https://motus.org/data/tagDeployment?id=38868 |
| 38873 | F | 2022-05-01 22:30 | N 52.247 E 1.619 | 2022-05-02 21:54 | N 52.247 E 1.615 | 2022-05-06 20:22 | N 51.759 E 3.845 | crossing in the night of 2, 3, 4 or 5 May (early evening detection 6 May in the Netherlands) | https://motus.org/data/tagDeployment?id=38873 |
| 35905 | F | 2022-05-01 23:45 | N 52.247 E 1.619 | 2022-05-06 20:07 | N 52.247 E 1.615 | 2022-05-09 22:21 | N 51.1156 E 2.638 | crossing was in the night of 6, 7 or 8 May (crossing in the night of 9 May highly unlikely, given the low groundspeed of 16 km/h) | https://motus.org/data/tagDeployment?id=35905 |
| 35901 | F | 2022-05-05 22:15 | N 51.938 E 1.320 | 2022-05-05 22:28 | N 51.938 E 1.320 | 2022-05-13 01:45 | N 53.300 E 5.078 | first detection in Netherlands late night 2022-05-13. Consequently, crossing was in the night of 5, 6, 7, 8, 9, 10, 11 or 12 May | https://motus.org/data/tagDeployment?id=35901 |
| 38998 | F | 2022-05-05 23:05 | N 51.938 E 1.320 | 2022-05-06 23:25 | N 51.938 E 1.320 | | | | https://motus.org/data/tagDeployment?id=38998 |
| 39000 | F | 2022-05-05 23:10 | N 51.938 E 1.320 | 2022-05-16 21:19 | N 52.656 E 1.729 | | | | https://motus.org/data/tagDeployment?id=39000 |
| 39001 | F | 2022-05-05 23:40 | N 51.938 E 1.320 | 2022-05-06 21:03 | N 51.938 E 1.320 | 2022-05-11 23:06 | N 53.630 E 9.589 | crossing in the night of 6, 7, 8, 9 or 10 May (given the evening detection in Germany on 11 May) | https://motus.org/data/tagDeployment?id=39001 |
| 39006 | M | 2022-05-10 21:40 | N 52.247 E 1.619 | NA | NA | | | tag detached 2022-05-17 | https://motus.org/data/tagDeployment?id=39006 |
| 39007 | F | 2022-05-11 02:10 | N 52.247 E 1.619 | 2022-05-11 21:10 | N 52.247 E 1.615 | 2022-05-12 00:00 | N 51.893 E 4.037 | crossing night 2022-05-11 | https://motus.org/data/tagDeployment?id=39007 |
| 45000 | M | 2023-03-30 20:00 | N 52.247 E 1.619 | 2023-04-29 20:16 | N 52.247 E 1.615 | | | | https://motus.org/data/tagDeployment?id=45000 |
| 42291 | M | 2023-04-28 21:30 | N 52.247 E 1.619 | 2023-05-27 23:46 | N 52.247 E 1.615 | | | | https://motus.org/data/tagDeployment?id=42291 |
| 42267 | F | 2023-04-28 22:00 | N 52.247 E 1.619 | 2023-05-28 02:10 | N 52.247 E 1.615 | | | | https://motus.org/data/tagDeployment?id=42267 |
| 42210 | F | 2023-04-28 23:15 | N 52.247 E 1.619 | 2023-04-30 21:17 | N 52.459 E 1.740 | | | | https://motus.org/data/tagDeployment?id=42210 |
| 43266 | F | 2023-04-29 00:15 | N 52.247 E 1.619 | 2023-05-07 23:56 | N 52.646 E 1.736 | | | | https://motus.org/data/tagDeployment?id=43266 |
| 43271 | F | 2023-04-29 00:30 | N 52.247 E 1.619 | 2023-04-29 20:50 | N 52.247 E 1.615 | | | | https://motus.org/data/tagDeployment?id=43271 |
| 43270 | F | 2023-04-29 01:00 | N 52.247 E 1.619 | 2023-04-29 20:01 | N 52.400 E 1.727 | | | | https://motus.org/data/tagDeployment?id=43270 |
| 43272 | F | 2023-04-29 01:15 | N 52.247 E 1.619 | 2023-05-04 22:08 | N 52.459 E 1.740 | 2023-05-05 23:14 | N 52.934 E 5.039 | crossing night 2023-05-04, as detected relatively early in the night in the Netherlands about 25 km inland on 5 May | https://motus.org/data/tagDeployment?id=43272 |
| 42325 | F | 2023-04-29 22:30 | N 52.387 E 1.707 | 2023-05-10 21:03 | N 52.459 E 1.740 | | | | https://motus.org/data/tagDeployment?id=42325 |
| 43051 | F | 2023-04-29 22:50 | N 52.387 E 1.707 | 2023-05-10 21:04 | N 52.646 E 1.736 | | | | https://motus.org/data/tagDeployment?id=43051 |

| Deployment | Sex | Tagged [UTC] | Lat/Long | Last UK [UTC] | Lat/Long | First mainland [UTC] | Lat/Long | Comments | Link |
|---|---|---|---|---|---|---|---|---|---|
| 43047 | F | 2023-04-30 00:15 | N 52.387 E 1.707 | 2023-04-30 00:25 | N 52.386 E 1.708 | 2023-05-08 22:28 | N 53.501 E 8.475 | crossing in the night of 1 - 7 May (30 April unlikely and 8 May not feasible, as detected relatively early in the evening in Germany) | https://motus.org/data/tagDeployment?id=43047 |
| 43054 | F | 2023-04-30 21:30 | N 52.247 E 1.619 | 2023-05-14 23:13 | N 52.801 E 1.583 | | | | https://motus.org/data/tagDeployment?id=43054 |
| 43053 | F | 2023-04-30 22:00 | N 52.247 E 1.619 | 2023-05-24 01:15 | N 52.247 E 1.615 | | | | https://motus.org/data/tagDeployment?id=43053 |
| 43273 | F | 2023-04-30 22:15 | N 52.247 E 1.619 | 2023-05-04 21:47 | N 51.938 E 1.320 | 2023-05-08 22:05 | N 53.437 E 8.092 | crossing in the night of 4, 5, 6 or 7 May | https://motus.org/data/tagDeployment?id=43273 |
| 42265 | F | 2023-04-30 22:45 | N 52.247 E 1.619 | 2023-05-09 20:50 | N 50.917 E 0.965 | | | this bat crossed from Dunwich to Koksijde in the night of 2023-05-01, and returned to the UK (Dungeness) the same night! | https://motus.org/data/tagDeployment?id=42265 |
| 43049 | M | 2023-04-30 23:15 | N 52.247 E 1.619 | 2023-05-10 21:09 | N 52.400 E 1.727 | 2023-05-11 20:01 | N 51.749 E 3.826 | crossing night 2023-05-10 | https://motus.org/data/tagDeployment?id=43049 |
| 43048 | F | 2023-04-30 23:45 | N 52.247 E 1.619 | 2023-05-04 21:55 | N 52.253 E 1.628 | | | | https://motus.org/data/tagDeployment?id=43048 |
| 43050 | M | 2023-05-01 00:15 | N 52.247 E 1.619 | 2023-05-04 20:50 | N 52.400 E 1.727 | | | | https://motus.org/data/tagDeployment?id=43050 |
| 43052 | F | 2023-05-01 00:45 | N 52.247 E 1.619 | 2023-05-14 21:30 | N 52.801 E 1.583 | | | | https://motus.org/data/tagDeployment?id=43052 |
| 43046 | F | 2023-05-01 00:50 | N 52.247 E 1.619 | 2023-05-14 21:25 | N 52.646 E 1.736 | 2023-05-15 02:24 | N 51.749 E 3.826 | crossing night 2023-05-14 | https://motus.org/data/tagDeployment?id=43046 |
| 42438 | F | 2023-05-01 01:15 | N 52.247 E 1.619 | 2023-05-04 22:45 | N 52.253 E 1.628 | 2023-05-28 00:41 | N 53.548 E 8.088 | crossing in the night between 4 - 27 May (given the detection in Germany in the night of 28 May) | https://motus.org/data/tagDeployment?id=42438 |
| 43055 | F | 2023-05-01 01:30 | N 52.247 E 1.619 | 2023-05-05 20:24 | N 52.801 E 1.583 | | | | https://motus.org/data/tagDeployment?id=43055 |
| 42440 | F | 2023-05-01 22:00 | N 52.247 E 1.619 | 2023-05-04 22:41 | N 52.646 E 1.736 | | | | https://motus.org/data/tagDeployment?id=42440 |
| 42442 | M | 2023-05-01 22:10 | N 52.247 E 1.619 | 2023-05-04 22:13 | N 52.400 E 1.727 | | | | https://motus.org/data/tagDeployment?id=42442 |
| 43058 | F | 2023-05-01 21:50 | N 52.247 E 1.619 | 2023-05-08 20:14 | N 52.253 E 1.628 | | | | https://motus.org/data/tagDeployment?id=43058 |
| 43056 | M | 2023-05-01 22:50 | N 52.247 E 1.619 | 2023-05-14 20:45 | N 52.386 E 1.708 | | | | https://motus.org/data/tagDeployment?id=43056 |
| 43057 | F | 2023-05-01 22:55 | N 52.247 E 1.619 | 2023-05-10 21:25 | N 52.386 E 1.708 | | | | https://motus.org/data/tagDeployment?id=43057 |
| 42237 | F | 2023-05-02 01:00 | N 52.247 E 1.619 | 2023-05-26 22:42 | N 52.801 E 1.583 | | | | https://motus.org/data/tagDeployment?id=42237 |
| 45001 | F | 2023-05-02 02:30 | N 52.247 E 1.619 | 2023-05-02 03:00 | N 52.247 E 1.615 | | | | https://motus.org/data/tagDeployment?id=45001 |
| 43269 | F | 2023-05-02 21:00 | N 52.387 E 1.707 | 2023-05-16 21:35 | N 52.253 E 1.628 | | | | https://motus.org/data/tagDeployment?id=43269 |
| 45002 | F | 2023-05-03 21:00 | N 52.247 E 1.619 | 2023-05-04 22:03 | N 52.400 E 1.727 | | | | https://motus.org/data/tagDeployment?id=45002 |
| 45003 | M | 2023-05-03 21:45 | N 52.247 E 1.619 | 2023-05-04 23:36 | N 52.459 E 1.740 | | | | https://motus.org/data/tagDeployment?id=45003 |
| 45004 | F | 2023-05-03 23:30 | N 52.247 E 1.619 | 2023-05-27 23:08 | N 52.646 E 1.736 | | | | https://motus.org/data/tagDeployment?id=45004 |
| 45005 | F | 2023-05-04 01:00 | N 52.247 E 1.619 | 2023-05-15 21:55 | N 52.253 E 1.628 | | | | https://motus.org/data/tagDeployment?id=45005 |
| 45006 | F | 2023-05-04 01:45 | N 52.247 E 1.619 | 2023-05-23 22:58 | N 52.253 E 1.628 | | | | https://motus.org/data/tagDeployment?id=45006 |
| 45008 | F | 2023-05-04 01:50 | N 52.247 E 1.619 | 2023-05-19 22:59 | N 51.599 E 0.936 | | | | https://motus.org/data/tagDeployment?id=45008 |
| 45009 | F | 2023-05-04 01:55 | N 52.247 E 1.619 | 2023-06-02 23:20 | N 51.938 E 1.320 | | | | https://motus.org/data/tagDeployment?id=45009 |
| 45010 | M | 2023-05-04 02:00 | N 52.247 E 1.619 | 2023-06-02 22:15 | N 52.253 E 1.628 | | | | https://motus.org/data/tagDeployment?id=45010 |
| 45011 | F | 2023-05-04 02:05 | N 52.247 E 1.619 | 2023-05-13 23:08 | N 52.400 E 1.727 | | | | https://motus.org/data/tagDeployment?id=45011 |
| 45012 | F | 2023-05-04 02:15 | N 52.247 E 1.619 | 2023-05-08 00:57 | N 52.801 E 1.583 | | | | https://motus.org/data/tagDeployment?id=45012 |
| 45013 | F | 2023-05-04 02:20 | N 52.247 E 1.619 | 2023-05-15 20:57 | N 52.253 E 1.628 | | | | https://motus.org/data/tagDeployment?id=45013 |
| 45014 | F | 2023-05-04 02:30 | N 52.247 E 1.619 | 2023-05-05 20:37 | N 52.247 E 1.615 | 2023-05-17 21:44 | N 51.489 E 4.057 | crossing in the night between 5 - 16 May (given the relatively early detection at Yerseke in the evening of 17 May) | https://motus.org/data/tagDeployment?id=45014 |
| 45016 | F | 2023-05-04 02:30 | N 52.247 E 1.619 | 2023-05-05 20:50 | N 52.253 E 1.628 | | | | https://motus.org/data/tagDeployment?id=45016 |
| 45017 | F | 2023-05-05 21:10 | N 52.387 E 1.707 | 2023-05-10 21:02 | N 52.386 E 1.708 | | | | https://motus.org/data/tagDeployment?id=45017 |
| 45018 | F | 2023-05-05 21:20 | N 52.387 E 1.707 | 2023-05-15 21:11 | N 52.646 E 1.736 | | | | https://motus.org/data/tagDeployment?id=45018 |
| 45019 | F | 2023-05-05 21:30 | N 52.387 E 1.707 | 2023-05-05 23:21 | N 52.386 E 1.708 | | | | https://motus.org/data/tagDeployment?id=45019 |
| 45020 | F | 2023-05-05 22:00 | N 52.387 E 1.707 | 2023-05-05 23:38 | N 52.459 E 1.740 | | | | https://motus.org/data/tagDeployment?id=45020 |
| 45021 | F | 2023-05-05 22:15 | N 52.387 E 1.707 | 2023-05-05 23:15 | N 52.386 E 1.708 | | | | https://motus.org/data/tagDeployment?id=45021 |
| 45022 | F | 2023-05-05 22:30 | N 52.387 E 1.707 | 2023-05-06 00:24 | N 52.646 E 1.736 | | | | https://motus.org/data/tagDeployment?id=45022 |
| 45023 | F | 2023-05-05 22:45 | N 52.387 E 1.707 | 2023-05-28 00:36 | N 51.938 E 1.320 | | | | https://motus.org/data/tagDeployment?id=45023 |
| 45024 | F | 2023-05-05 22:50 | N 52.387 E 1.707 | 2023-05-17 01:44 | N 52.400 E 1.727 | | | | https://motus.org/data/tagDeployment?id=45024 |
| 45025 | F | 2023-05-06 00:30 | N 52.387 E 1.707 | 2023-05-08 22:20 | N 52.253 E 1.628 | | | | https://motus.org/data/tagDeployment?id=45025 |
| 45026 | F | 2023-05-07 20:45 | N 52.247 E 1.619 | 2023-05-15 20:35 | N 52.253 E 1.628 | | | | https://motus.org/data/tagDeployment?id=45026 |
| 45027 | F | 2023-05-07 21:45 | N 52.247 E 1.619 | 2023-05-23 23:35 | N 52.646 E 1.736 | | | | https://motus.org/data/tagDeployment?id=45027 |
| 45028 | F | 2023-05-07 21:50 | N 52.247 E 1.619 | 2023-05-21 21:12 | N 52.253 E 1.628 | 2023-05-25 00:13 | N 51.358 E 3.349 | crossing in the night of 21, 22, 23 or 24 May | https://motus.org/data/tagDeployment?id=45028 |
| 45029 | F | 2023-05-07 22:10 | N 52.247 E 1.619 | 2023-06-02 22:05 | N 51.938 E 1.320 | | | | https://motus.org/data/tagDeployment?id=45029 |
| 44980 | F | 2023-05-07 22:30 | N 52.247 E 1.619 | 2023-05-15 20:52 | N 52.253 E 1.628 | | | | https://motus.org/data/tagDeployment?id=44980 |
| 44981 | M | 2023-05-07 22:45 | N 52.247 E 1.619 | 2023-06-04 01:34 | N 52.253 E 1.628 | | | | https://motus.org/data/tagDeployment?id=44981 |
| 44982 | F | 2023-05-07 22:50 | N 52.247 E 1.619 | 2023-05-20 02:22 | N 52.247 E 1.615 | | | | https://motus.org/data/tagDeployment?id=44982 |
| 44984 | F | 2023-05-07 22:55 | N 52.247 E 1.619 | 2023-05-10 21:08 | N 52.253 E 1.628 | 2023-05-14 22:34 | N 51.124 E 2.648 | crossing in the night of 10, 11, 12, 13 or 14 May | https://motus.org/data/tagDeployment?id=44984 |
| 44983 | M | 2023-05-07 23:00 | N 52.247 E 1.619 | 2023-05-19 20:49 | N 52.253 E 1.628 | | | | https://motus.org/data/tagDeployment?id=44983 |
| 44988 | F | 2023-05-07 23:10 | N 52.247 E 1.619 | 2023-05-08 00:40 | N 52.247 E 1.615 | | | | https://motus.org/data/tagDeployment?id=44988 |
| 44987 | F | 2023-05-07 23:20 | N 52.247 E 1.619 | 2023-05-22 20:44 | N 52.253 E 1.628 | | | | https://motus.org/data/tagDeployment?id=44987 |
| 44986 | F | 2023-05-07 23:30 | N 52.247 E 1.619 | 2023-05-15 22:05 | N 50.917 E 0.965 | | | | https://motus.org/data/tagDeployment?id=44986 |
| 44989 | F | 2023-05-08 00:40 | N 52.247 E 1.619 | 2023-05-14 23:06 | N 51.938 E 1.320 | | | | https://motus.org/data/tagDeployment?id=44989 |
| 44990 | F | 2023-05-08 01:10 | N 52.247 E 1.619 | 2023-05-08 01:35 | N 52.247 E 1.615 | | | | https://motus.org/data/tagDeployment?id=44990 |
| 44991 | F | 2023-05-08 01:20 | N 52.247 E 1.619 | 2023-05-11 02:25 | N 51.2678 E 1.3743 | | | | https://motus.org/data/tagDeployment?id=44991 |
| 44992 | F | 2023-05-08 02:00 | N 52.247 E 1.619 | 2023-05-14 21:11 | N 52.247 E 1.615 | | | | https://motus.org/data/tagDeployment?id=44992 |
| 44993 | F | 2023-05-08 02:40 | N 52.247 E 1.619 | 2023-05-08 21:05 | N 52.253 E 1.628 | | | | https://motus.org/data/tagDeployment?id=44993 |
| 44995 | F | 2023-05-10 22:45 | N 52.247 E 1.619 | 2023-05-20 21:47 | N 52.253 E 1.628 | | | | https://motus.org/data/tagDeployment?id=44995 |
| 44996 | F | 2023-05-10 22:50 | N 52.247 E 1.619 | 2023-05-21 22:49 | N 52.386 E 1.708 | | | | https://motus.org/data/tagDeployment?id=44996 |
| 45007 | M | 2023-05-10 22:50 | N 52.247 E 1.619 | 2023-06-11 00:25 | N 51.2678 E 1.3743 | 2023-06-15 00:27 | N 51.622 E 3.669 | crossing in the night of 11, 12, 13, 14 or 15 June | https://motus.org/data/tagDeployment?id=45007 |
| 44994 | F | 2023-05-11 01:00 | N 52.247 E 1.619 | 2023-05-14 20:58 | N 52.253 E 1.628 | 2023-05-15 01:36 | N 51.983 E 4.116 | crossing night 2023-05-14 | https://motus.org/data/tagDeployment?id=44994 |
| 44997 | F | 2023-05-11 01:40 | N 52.247 E 1.619 | 2023-05-16 21:21 | N 52.400 E 1.727 | | | | https://motus.org/data/tagDeployment?id=44997 |
| 44998 | F | 2023-05-11 02:10 | N 52.247 E 1.619 | 2023-05-16 20:22 | N 52.253 E 1.628 | 2023-05-17 01:02 | N 51.529 E 3.447 | crossing night 2023-05-16 | https://motus.org/data/tagDeployment?id=44998 |
| 44999 | F | 2023-05-11 02:40 | N 52.247 E 1.619 | 2023-05-17 22:32 | N 52.646 E 1.736 | | | | https://motus.org/data/tagDeployment?id=44999 |
| 45015 | M | 2023-05-11 02:45 | N 52.247 E 1.619 | 2023-05-14 23:18 | N 52.459 E 1.740 | | | | https://motus.org/data/tagDeployment?id=45015 |
| 44985 | M | 2023-05-11 02:45 | N 52.247 E 1.619 | 2023-06-04 00:42 | N 52.253 E 1.628 | | | | https://motus.org/data/tagDeployment?id=44985 |

A total of 27 bats was detected on the European mainland, excluding deployment 42265. This individual crossed the North Sea in the night of 1 May 2023 twice: from Dunwich (UK) to Koksijde in Belgium (min. 144 km) and back to Dungeness (UK) (min. 120 km). Figure SF1-1 shows the movements between the last detection in the UK and the first detection on the European mainland. Note that the shortest distance between the receivers is shown, which not necessarily reflects the actual flight path between the receivers.

*Figure SF 1-1: Movements of Nathusius' pipistrelles included in the analysis (n=27). MOTUS receivers are indicated as yellow dots, tagging locations in red. Solid lines represent movements occurring within a single night, while dotted lines indicate movements over multiple nights.*

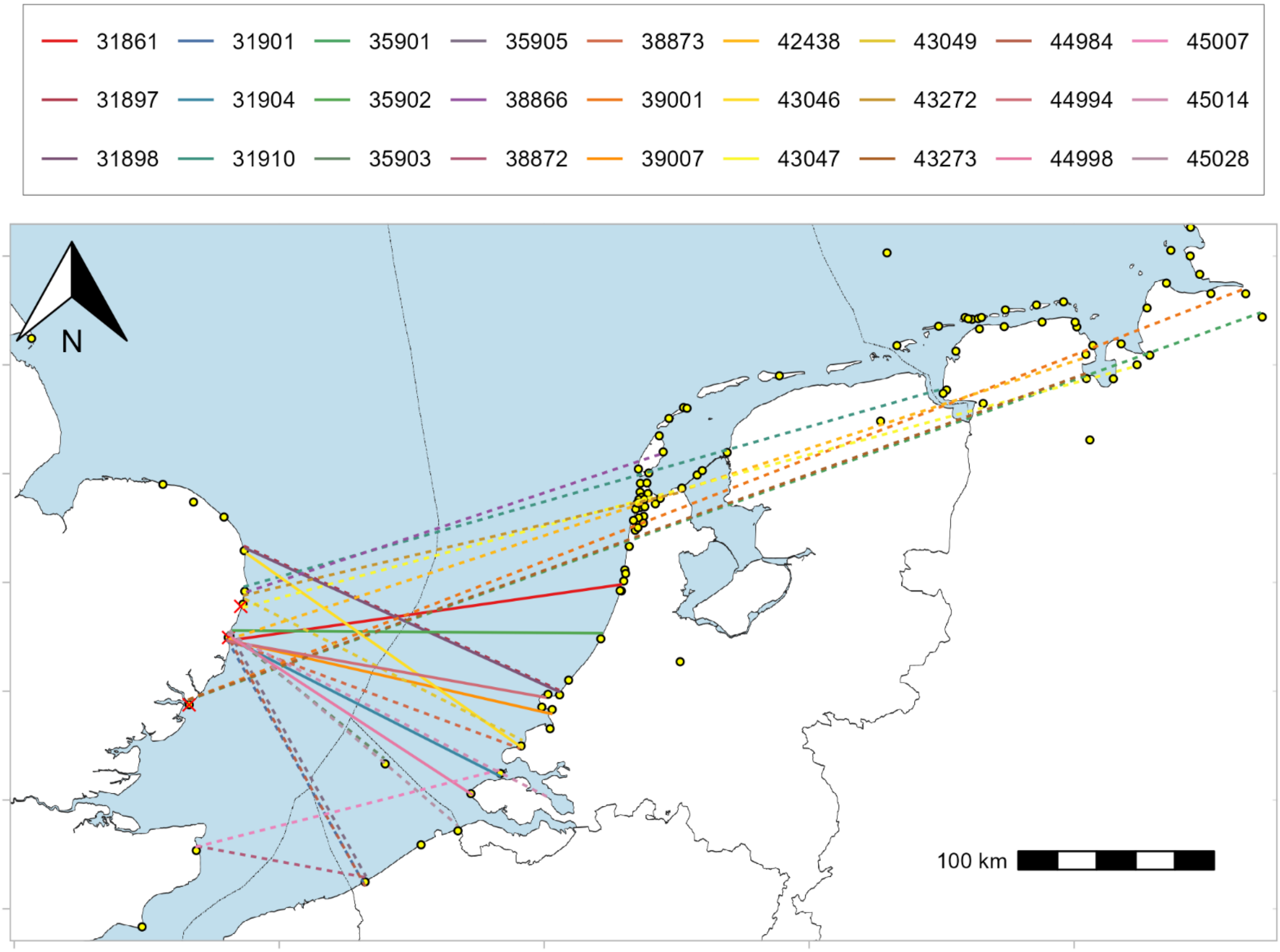
