## Supplementary file 2 for "Directional variation and method-specific detection patterns in offshore bat migration: implications for wind farm mitigation"

**Supplementary file 2: Acoustic monitoring equipment, monitoring locations and raw data processing**

*Equipment*

Acoustic bat activity was monitored with an Avisoft - UltraSoundGate 116Hnbm in combination with a omnidirectional electret ultrasound microphone FG-DT50 (Avisoft Bioacoustics, Germany). The settings of the UltraSoundGate recording software (RECORDER v. 4.2.29) are shown in Table SF2-1.

*Table SF2-1 Software settings of the UltraSoundGate*

| Parameter | Value | Parameter | Value |
|---|---|---|---|
| Pre-trigger | 0.1 s | Buffer | 0.064 s |
| Hold tm | 0.8 s | Bat call filter | Enabled |
| Duration | > 0 s | Accept monotonic structures | Enabled |
| Syllable | > 0 s | Min sweep rate FM | -20 KHz/ms |
| Reject wind/rain | enabled | Min sweep rate CF | -3 KHz/ms |
| Trigger event level | 0.501% | Max sweep rate FM | -1 KHz/ms |
| Trigger event range | 15-100kHz | Max sweep rate CF | 2 KHz/ms |
| Sampling rate | 250000Hz | Min duration FM | 1 ms |
| Format | 16 bit | Min duration CF | 2 ms |

The horizontal-mounted microphone was enclosed in a waterproof box (figure SF2-1) and connected with the soundgate through a S/FTP Cat 7 Marine Approved network cable at the monitoring locations Petrogas P9-A (Horizon) and Petrogas Q1-A (Helder). At all other monitoring locations a 2Triple x 0.75mm2 (CI2 TCC) RFOU(I)S1/S6 EPR/ICM/ZHAL/TCWB/ZHAL BLUE 250v cable was used.

The performance of the equipment was checked once per week at the monitoring locations with internet connectivity (C-Power, Belwind, Lichteiland Goeree, Dana P11-B, Luchterduinen,  Petrogas P9-A, PAWP, Petrogas Q1-A, Neptune K12-B & Neptune L10A-AC). The performance of the equipment at locations without internet connection (Europlatform, Wintershall P6-A & Wintershall K13-A) could not be checked regularly. At all locations, the microphones were replaced just before the spring migration season (late February/early March), and subsequently recalibrated by the manufacturer (Avisoft Bioacoustics).

*Monitoring locations*

We monitored at four offshore high voltage stations (OHVS) in wind farms (C-Power, Belwind, Luchterduinen, and PAWP), two measurement platforms (Europlatform and Lichteiland Goeree) and seven gas production platforms (Dana P11-B, Petrogas P9-A, Wintershall P6-A, Petrogas Q1-A, Wintershall K13-A, Neptune K12-BP and Neptune L10A-AC). The microphone was positioned at heights between 15 and 33 meters above sea level (mean height 21 meters). If technically feasible, the microphone was orientated in an easterly direction to minimize salt spray during strong westerlies (table SF2-2).

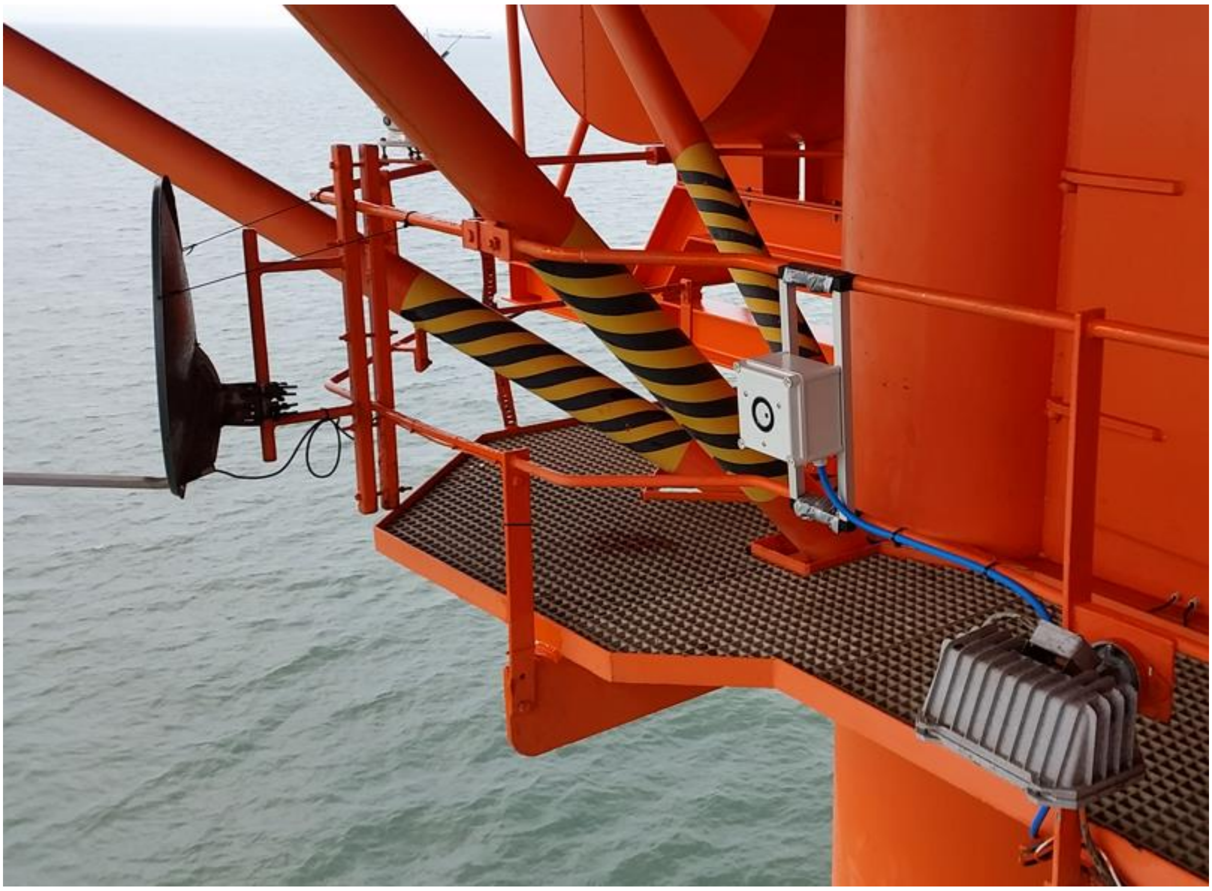

*Figure SF2-1: Acoustic monitoring at Lichteiland Goeree*

*Table SF2-2 Geographical location of the monitoring locations, height and orientation of the microphones*

| **Monitoring location** | **Longitude** | **Latitude** | **Height above sea level [m]** | **Orientation microphone [degrees from N]** |
|---|---|---|---|---|
| C-power OHVS | 2.99 | 51.57 | 15 | 60 |
| Belwind OHVS | 2.81 | 51.69 | 20 | 90 |
| Europlatform | 3.27 | 51.99 | 15 | 90 |
| Lichteiland Goeree | 3.66 | 51.92 | 15 | 90 |
| Dana P11-B | 3.34 | 52.35 | 25 | 90 |
| Luchterduinen OHVS | 4.17 | 52.40 | 15 | 90 |
| Petrogas P9-A (Horizon) | 3.74 | 52.55 | 33 | 45 |
| PAWP OHVS | 4.23 | 52.58 | 15 | 90 |
| Wintershall P6-A | 3.75 | 52.75 | 23 | 110 |
| Petrogas Q1-A (Helder) | 4.09 | 52.92 | 25 | 200 |
| Wintershall K13-A | 3.22 | 53.05 | 25 | 130 |
| Neptune K12-BP | 3.89 | 53.34 | 20 | 135 |
| Neptune L10A-AC | 4.20 | 53.40 | 17 | 90 |

Monitoring was executed from March until June during four consecutive years (2018-2021). In some cases downtime occurred due to malfunctioning equipment or logistical problems. The effective monitoring periods are shown in table SF2-3.

*Table SF2-3 Effective monitoring periods per location per year*

| **Monitoring location** | **2018** | **2019** | **2020** | **2021** |
|---|---|---|---|---|
| C-power OHVS | 01/03 - 30/06 | 01/03 - 30/06 | 21/06 - 30/06 | 01/03 - 30/06 |
| Belwind OHVS | 01/03 - 30/06 | 01/03 - 30/06 | 01/03 - 30/06 | - |
| Europlatform | 06/03 - 30/06 | 01/03 - 30/06 | 01/03 - 17/05 | 26/05 - 30/06 |
| Lichteiland Goeree | 06/03 - 30/06 | 09/05 - 30/06 | 24/06 - 30/06 | 03/03 - 10/03<br>22/04 - 02/05 |
| Dana P11-B | - | 01/03 - 27/06 | 01/03 - 06/06 | 01/03 - 30/06 |
| Luchterduinen OHVS | 04/04 - 30/06 | 01/03 - 30/06 | 06/04 - 12/06 | 05/03 - 07/05 |
| Petrogas P9-A (Horizon) | - | 01/03 - 30/06 | 01/03 - 08/06 | 06/03 - 30/06 |
| PAWP OHVS | 01/03 - 30/06 | 01/03 - 30/06 | 04/03 - 30/06 | 01/03 - 30/06 |
| Wintershall P6-A | 18/03 - 30/06 | 01/03 - 30/06 | 01/03 - 04/06 | 16/03 - 30/06 |
| Petrogas Q1-A (Helder) | - | 01/03 - 30/06 | 01/03 - 30/06 | 01/03 - 12/05 |
| Wintershall K13-A | - | 01/03 - 30/06 | - | - |
| Neptune K12-BP | 01/03 - 30/06 | 01/03 - 30/06 | 18/03 - 31/05 | 20/03 - 17/06 |
| Neptune L10A-AC | 01/03 - 30/06 | 01/03 - 30/06 | 21/03 - 08/04 | - |

*Processing of raw acoustic data*

Bats use echolocation to capture prey and to get an image of their environment. Wind gusts or maintenance and production activities at offshore platforms, however, also produce ultrasonic sounds. The first step in post-processing raw acoustic data is the separation of recordings with bat calls from recordings with other sound sources. We extracted bat call recordings from the raw acoustic data with the cross-correlation function of Avisoft SASlab Pro, version 5.2.14 (Avisoft bioacoustics), using recordings of reference calls of Nathusius' pipistrelle (n = 8), Common pipistrelle P. pipistrellus (n = 4), Pond bat Myotis dasycneme (n = 4), Common noctule Nyctalus noctula (n = 18), Serotine bat Eptesicus serotinus (n = 5) and the Nyctaloid group (n = 31, including the genera Nyctalus, Vespertilio, Eptesicus). Subsequently, each bat call was individually assessed and identified to the lowest possible taxonomic level, according to the criteria of (Barataud, 2020). Using visual inspection of the spectrograms of the recordings, feeding buzzes/intense exploration echolocation calls were identified (cf. Barataud, 2020), and subsequently verified by listening to the recordings.
