## Supplementary file 3 for "Directional variation and method-specific detection patterns in offshore bat migration: implications for wind farm mitigation"

**Supplementary file 3: Acoustic bat activity**

Figure SF3-1 shows the bat occurrence of the 172 distinguished individuals for all monitoring locations throughout the season and the night for each monitoring year.

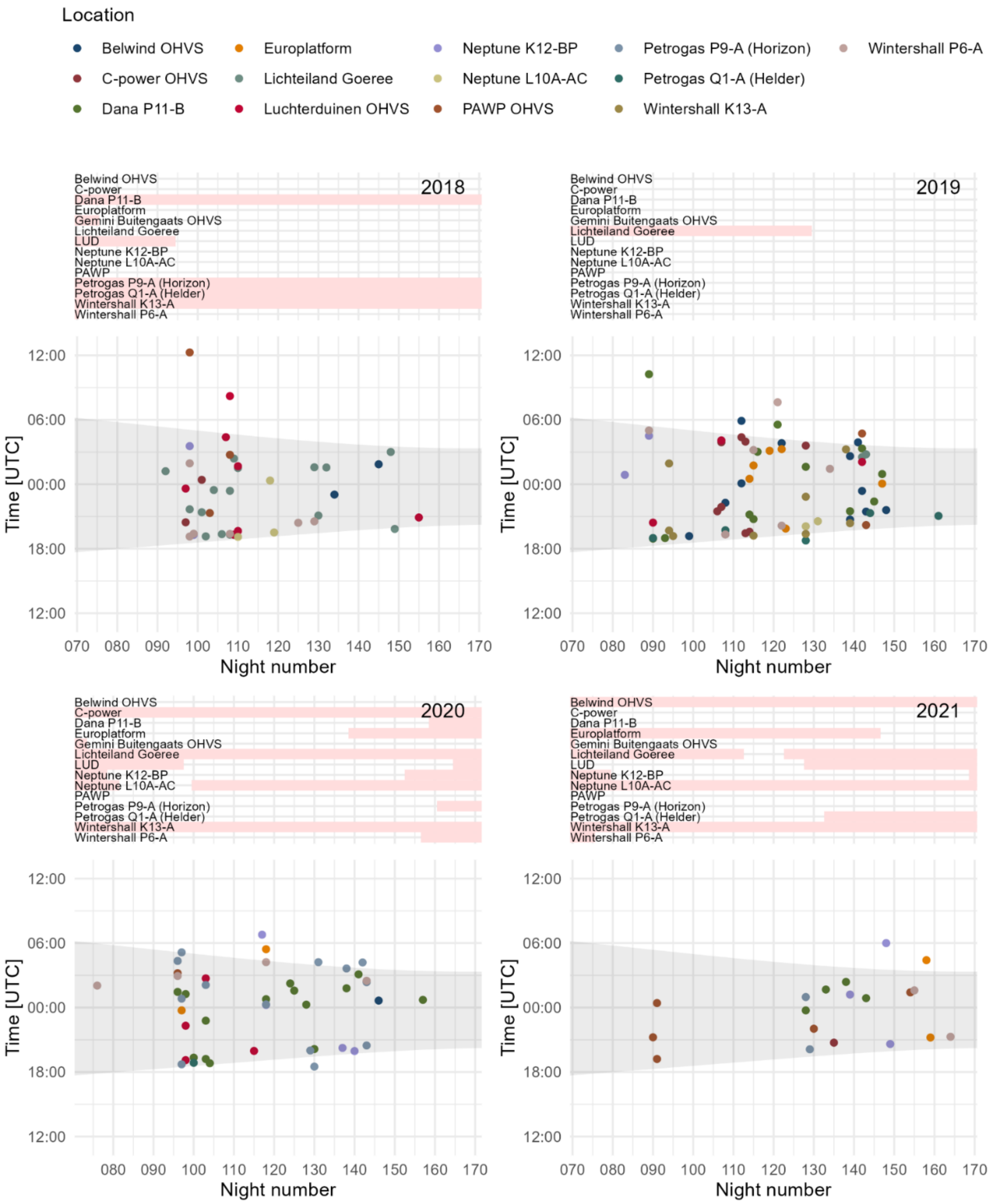


*Figure SF3-1: Bat occurrence throughout the night (time interval between sunset and sunrise is represented by grey) and the season for all monitoring locations (2018-2021). The effective monitoring periods per location are indicated in the headers with a white background, whereas the pink background indicates no monitoring. The dots represent the timestamp of the first recording of each individual.*

Figure SF3-2 shows the bat occurrence in the first half of the night (until midnight) in relation to the sunset and figure SF3-3 shows the occurrence in the second half of the night (after midnight) in relation to the sunrise. The timestamp of the first recording of each individual is used to determine the timeframe between sunset and sunrise respectively.

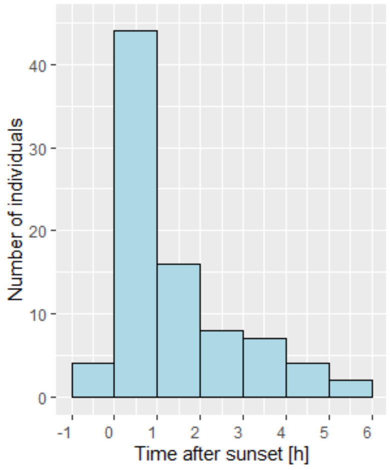


*Figure SF3-2: Histogram of bat occurrence before midnight [UTC] in relation to sunset, using the timestamp of the first recording for each individual*

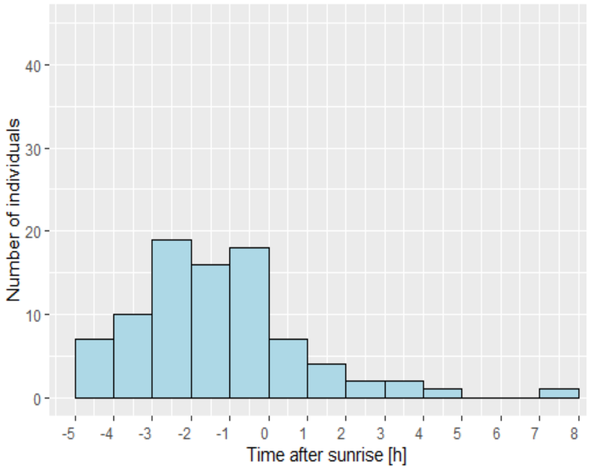


*Figure SF3-3: Histogram of bat occurrence after midnight [UTC] in relation to sunrise, using the timestamp of the first recording for each individual*

Figure SF3-4 shows the staging times of the distinguished individual bats.

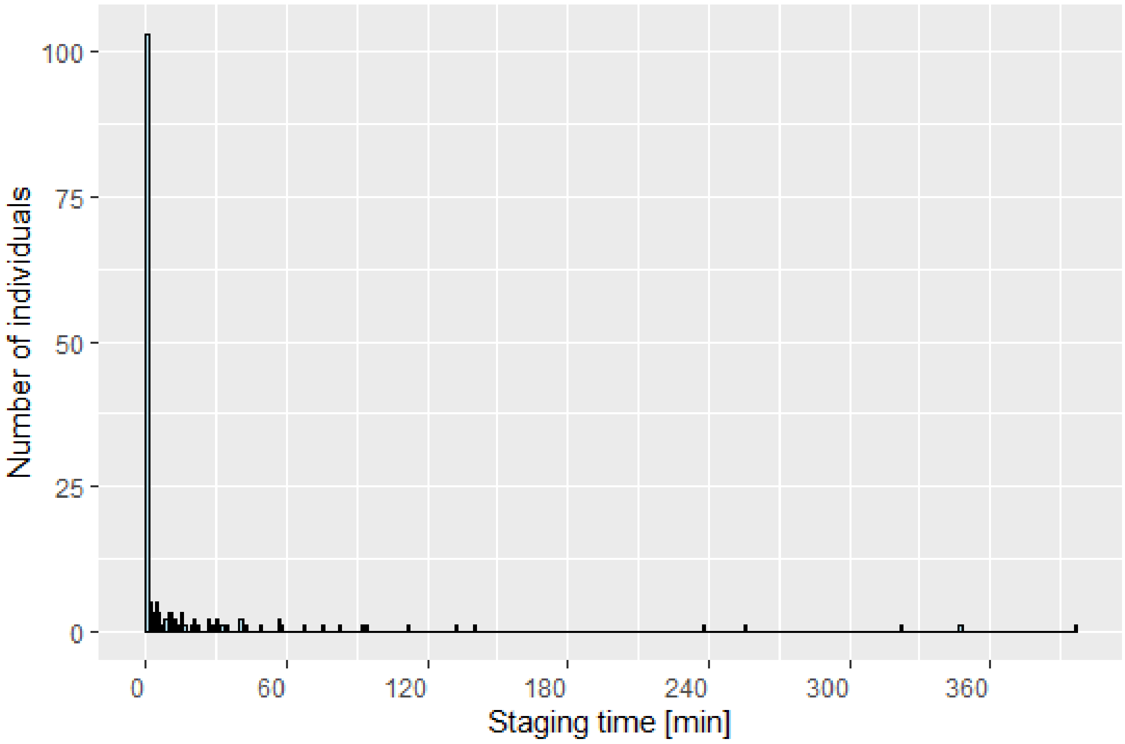


*Figure SF3-4: Histogram of staging times*

Figures SF3-5 – SF3-48 show the recorded acoustic presence of Nathusius' pipistrelle per monitoring location for each year. A total of 2777 Nathusius' pipistrelle recordings were obtained of which 152 (5.5%) contained feeding buzzes/intense exploration calls. Blue dots refer to recordings with exclusively echolocation calls, while red dots represent recordings with one or more feeding buzzes/intense exploration echolocation calls. The effective monitoring period is indicated by a white background, whereas a pink background indicates no monitoring. The time interval between sunset and sunrise is represented by grey. See supplementary file 1 for details on the monitoring locations.

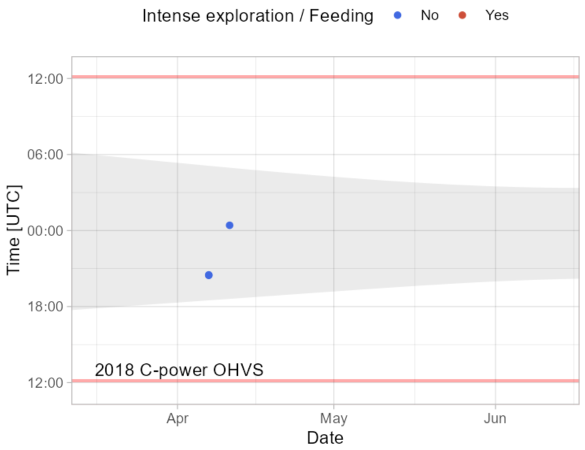


*Figure SF3-5: Recorded acoustic presence of Nathusius' pipistrelle - spring 2018 at C-Power OHVS*

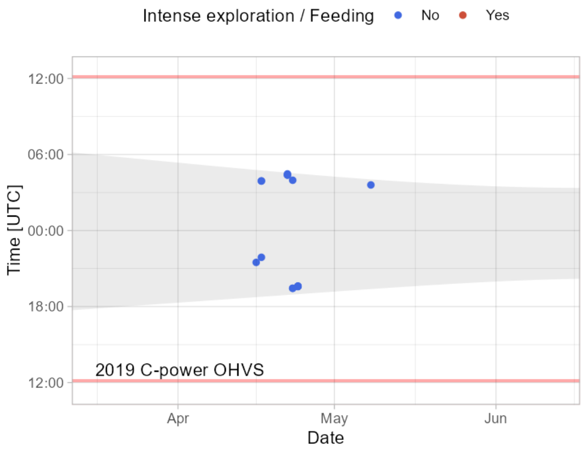


*Figure SF3-6: Recorded acoustic presence of Nathusius' pipistrelle - spring 2019 at C-Power OHVS*

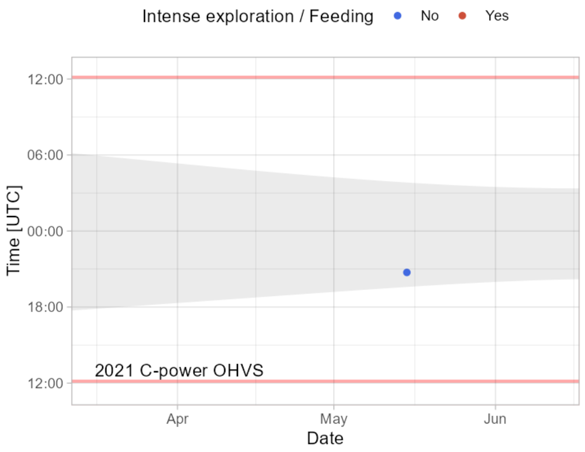


*Figure SF3-7: Recorded acoustic presence of Nathusius' pipistrelle - spring 2021 at C-Power OHVS*

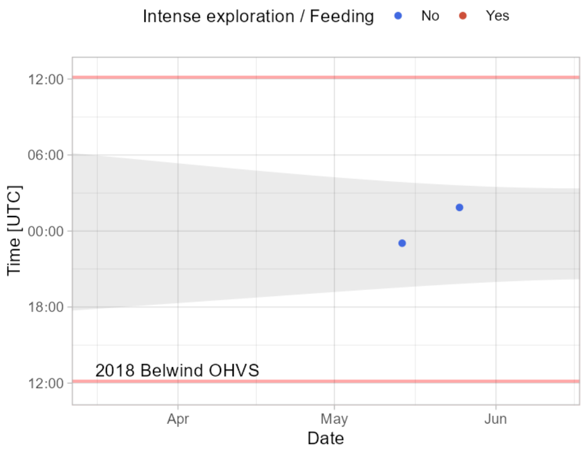


*Figure SF3-8: Recorded acoustic presence of Nathusius' pipistrelle - spring 2018 at Belwind OHVS*

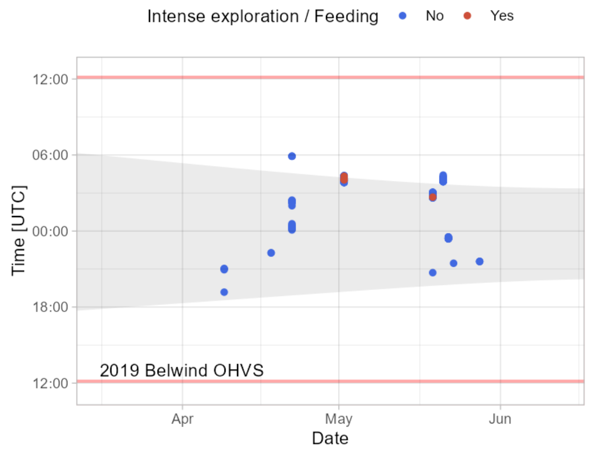


*Figure SF3-9: Recorded acoustic presence of Nathusius' pipistrelle - spring 2019 at Belwind OHVS*

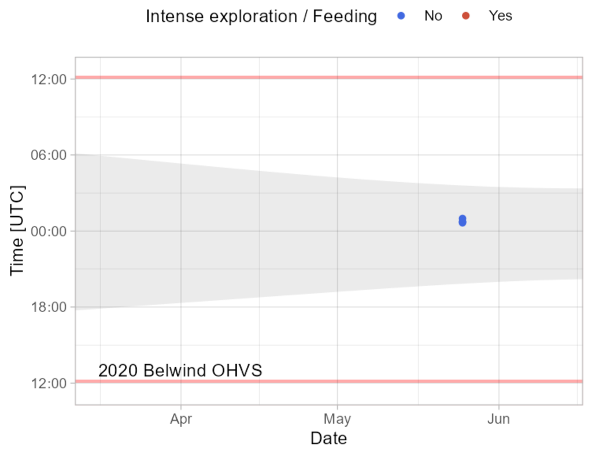


*Figure SF3-10: Recorded acoustic presence of Nathusius' pipistrelle - spring 2020 at Belwind OHVS*

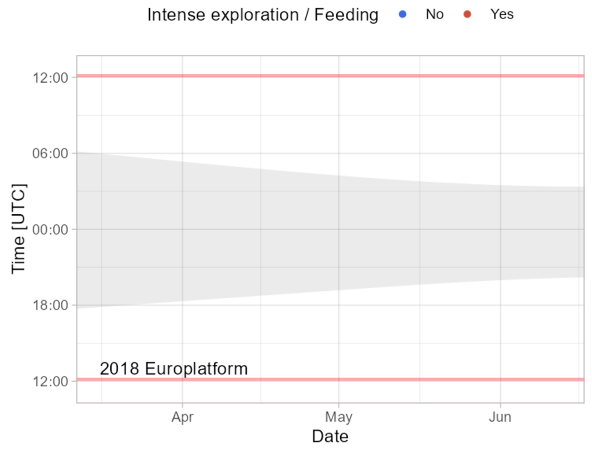


*Figure SF3-11: Recorded acoustic presence of Nathusius' pipistrelle - spring 2018 at Europlatform*

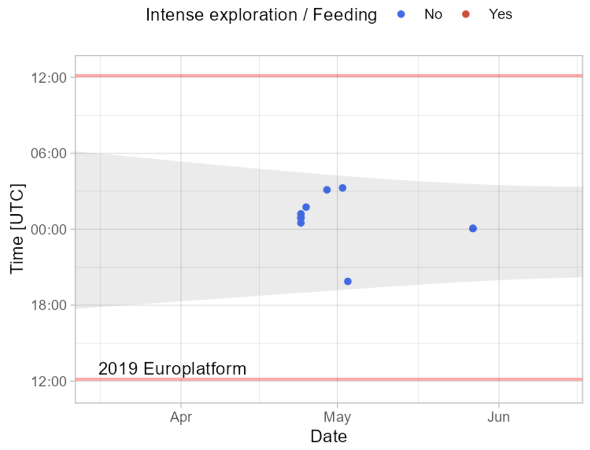


*Figure SF3-12: Recorded acoustic presence of Nathusius' pipistrelle – spring 2019 at Europlatform*

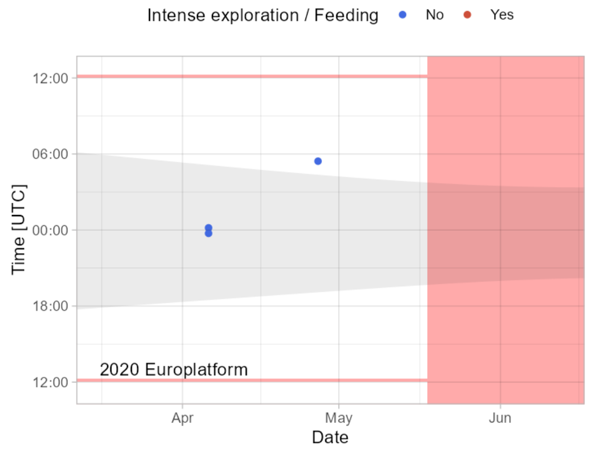


*Figure SF3-13: Recorded acoustic presence of Nathusius' pipistrelle – spring 2020 at Europlatform*

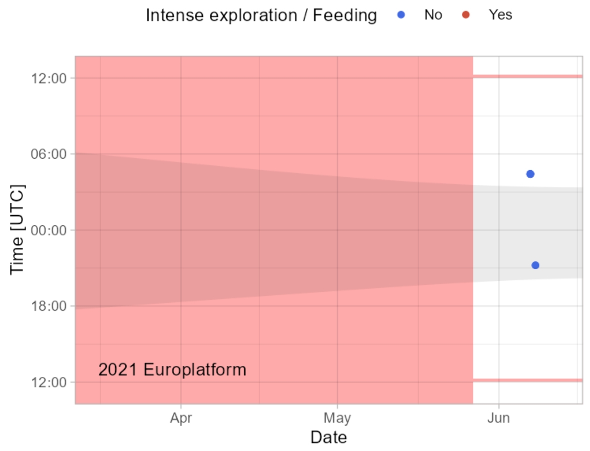


*Figure SF3-14: Recorded acoustic presence of Nathusius' pipistrelle – spring 2021 at Europlatform*

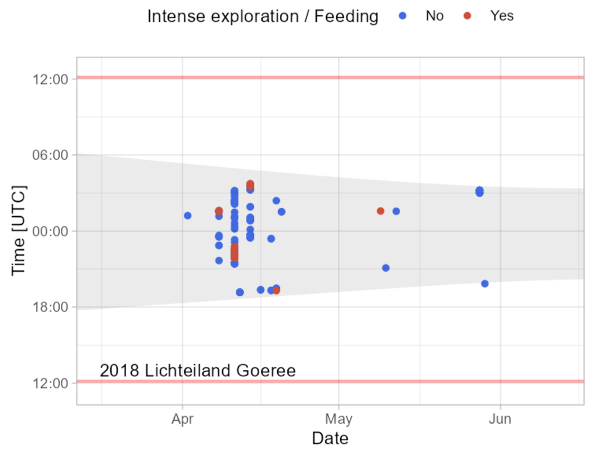


*Figure SF3-15: Recorded acoustic presence of Nathusius' pipistrelle – spring 2018 at Lichteiland Goeree*

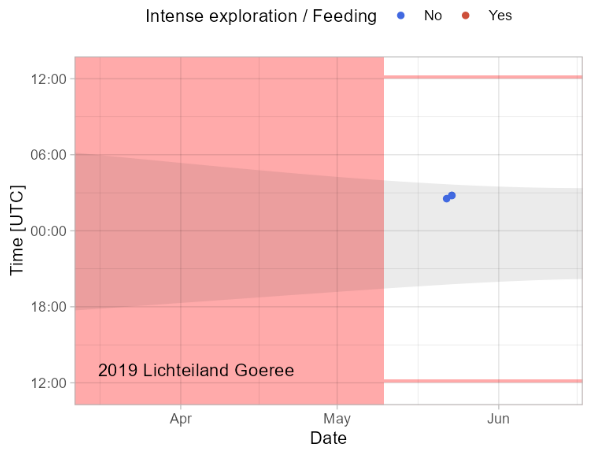


*Figure SF3-16: Recorded acoustic presence of Nathusius' pipistrelle – spring 2019 at Lichteiland Goeree*

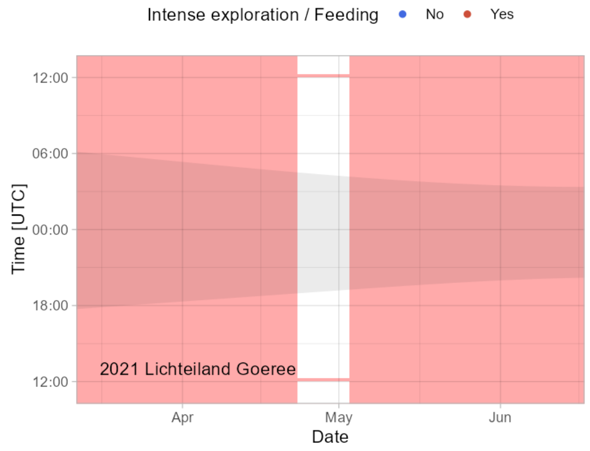


*Figure SF3-17: Recorded acoustic presence of Nathusius' pipistrelle – spring 2021 at Lichteiland Goeree*

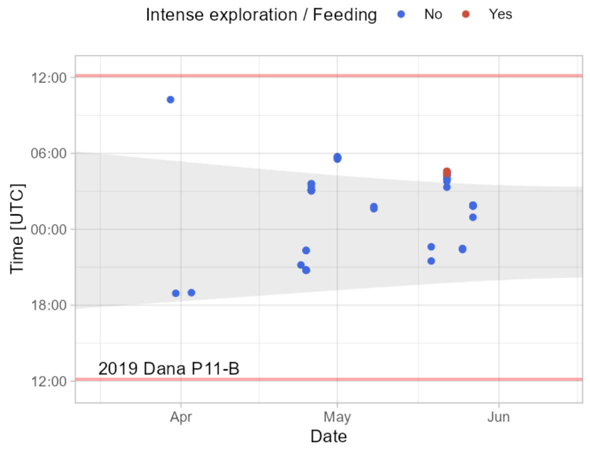


*Figure SF3-18: Recorded acoustic presence of Nathusius' pipistrelle – spring 2019 at Dana P11-B*

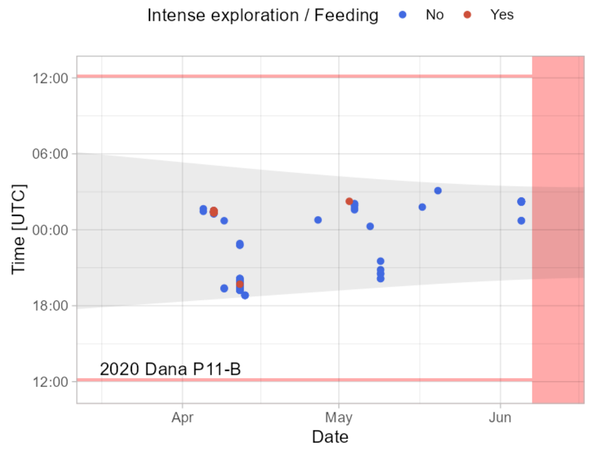


*Figure SF3-19: Recorded acoustic presence of Nathusius' pipistrelle – spring 2020 at Dana P11-B*

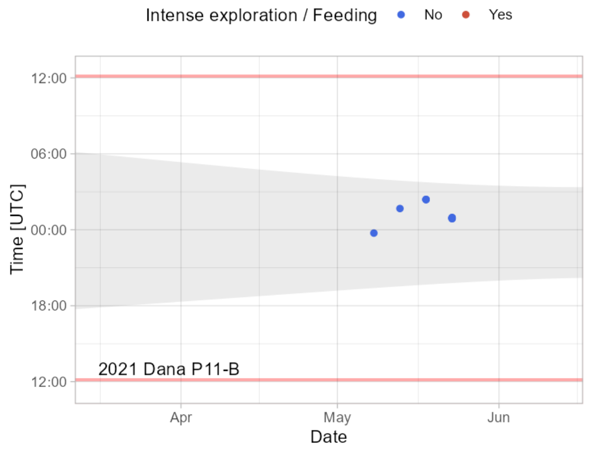


*Figure SF3-20: Recorded acoustic presence of Nathusius' pipistrelle – spring 2021 at Dana P11-B*

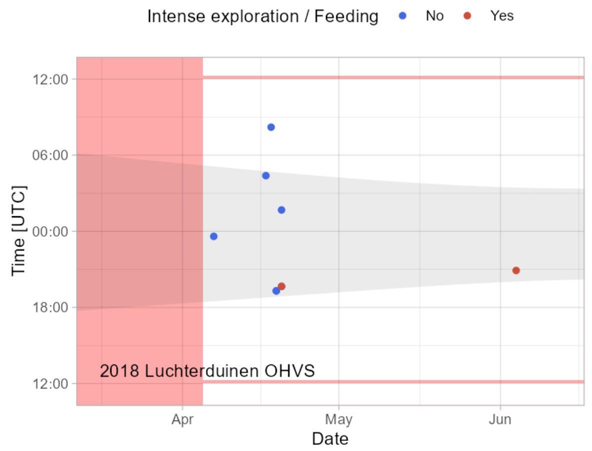


*Figure SF3-21: Recorded acoustic presence of Nathusius' pipistrelle – spring 2018 at Luchterduinen OHVS*

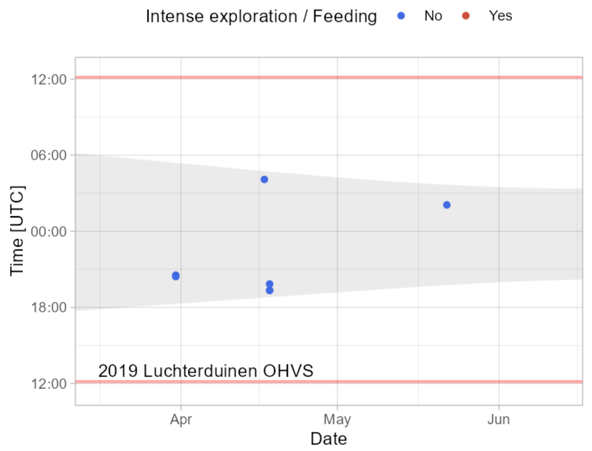


*Figure SF3-22: Recorded acoustic presence of Nathusius' pipistrelle – spring 2019 at Luchterduinen OHVS*

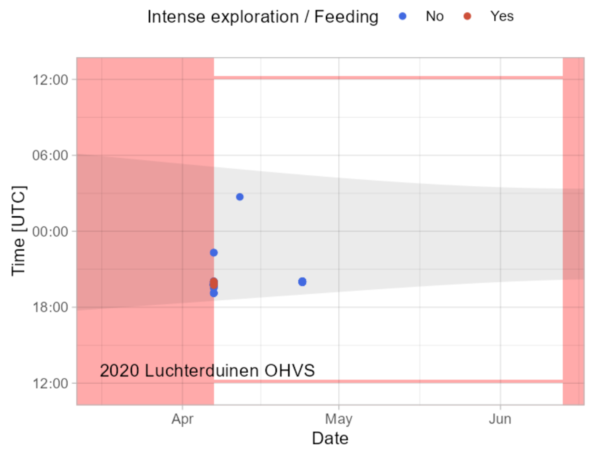


*Figure SF3-23: Recorded acoustic presence of Nathusius' pipistrelle – spring 2020 at Luchterduinen OHVS*

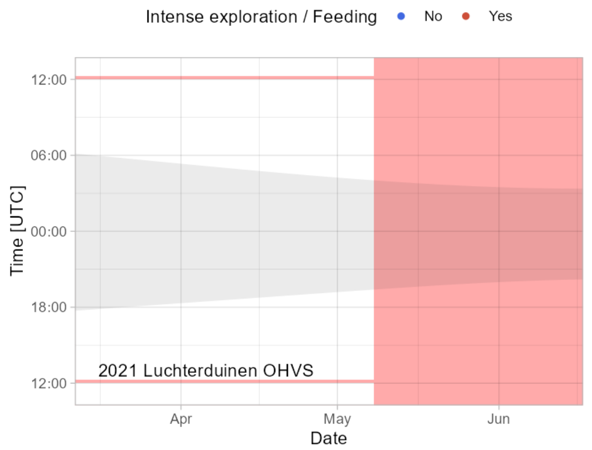


*Figure SF3-24: Recorded acoustic presence of Nathusius' pipistrelle – spring 2021 at Luchterduinen OHVS*

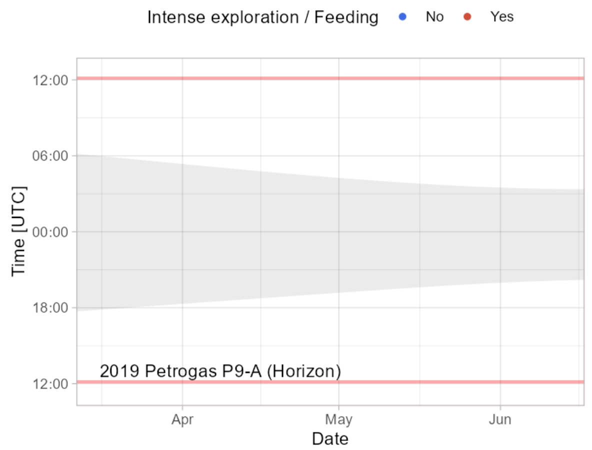


*Figure SF3-25: Recorded acoustic presence of Nathusius' pipistrelle – spring 2019 at Petrogas P9-A (Horizon)*

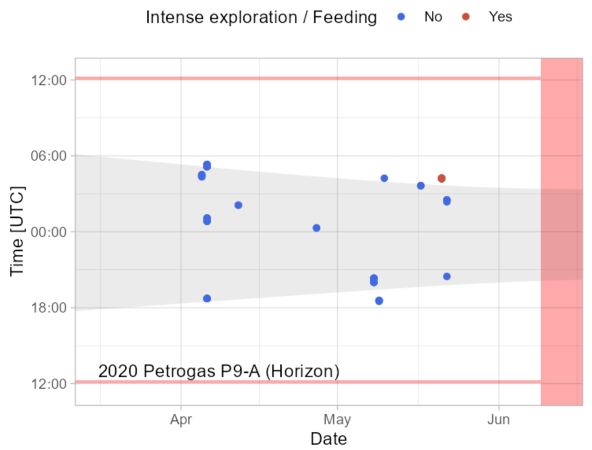


*Figure SF3-26: Recorded acoustic presence of Nathusius' pipistrelle – spring 2020 at Petrogas P9-A (Horizon)*

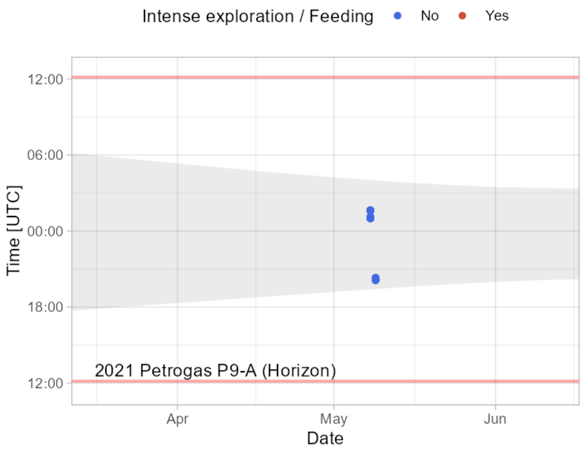


*Figure SF3-27: Recorded acoustic presence of Nathusius' pipistrelle – spring 2021 at Petrogas P9-A (Horizon)*

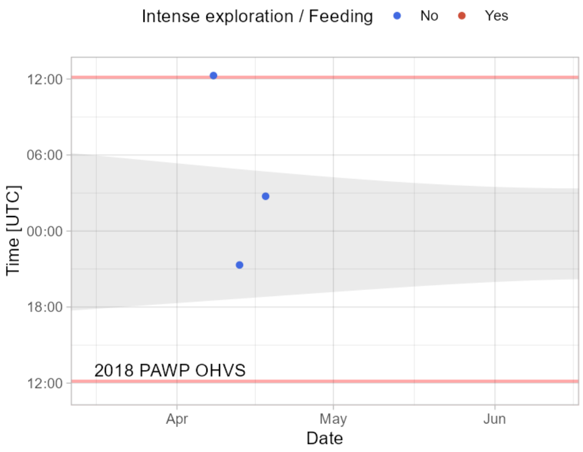


*Figure SF3-28: Recorded acoustic presence of Nathusius' pipistrelle – spring 2018 at PAWP*

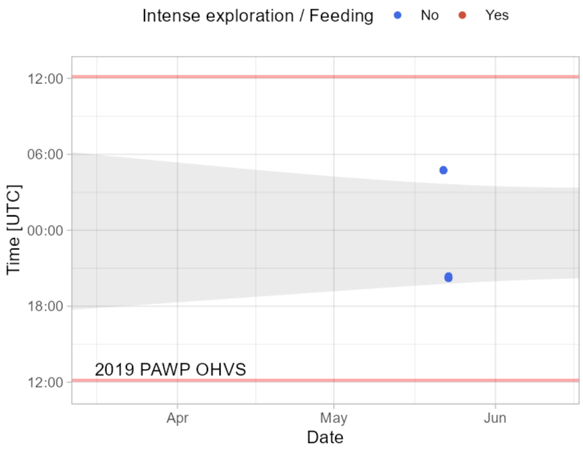


*Figure SF3-29: Recorded acoustic presence of Nathusius' pipistrelle – spring 2019 at PAWP*

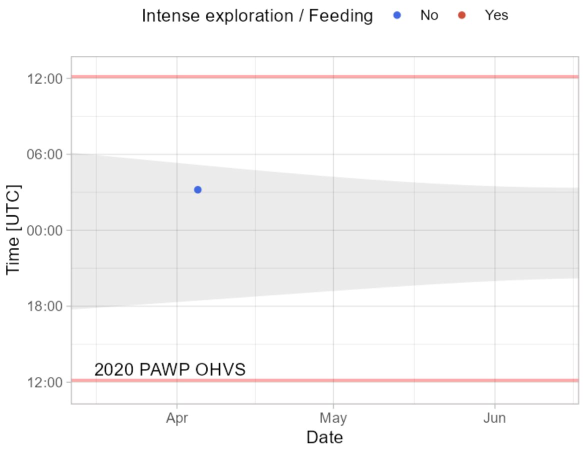


*Figure SF3-30: Recorded acoustic presence of Nathusius' pipistrelle – spring 2020 at PAWP*

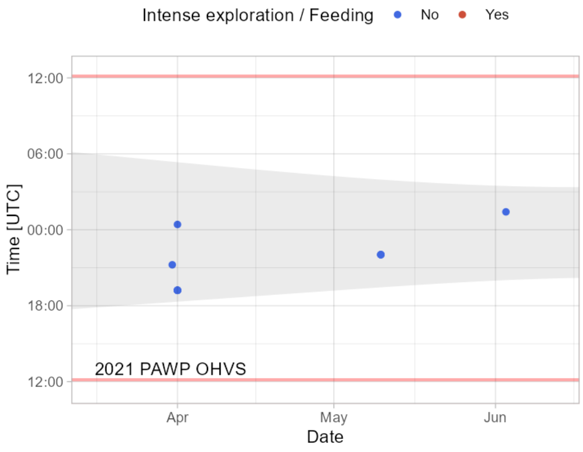


*Figure SF3-31: Recorded acoustic presence of Nathusius' pipistrelle – spring 2021 at PAWP*

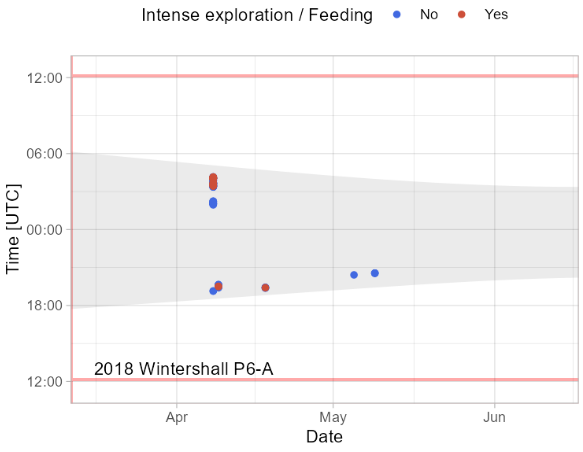


*Figure SF3-32: Recorded acoustic presence of Nathusius' pipistrelle – spring 2018 at Wintershall P6-A*

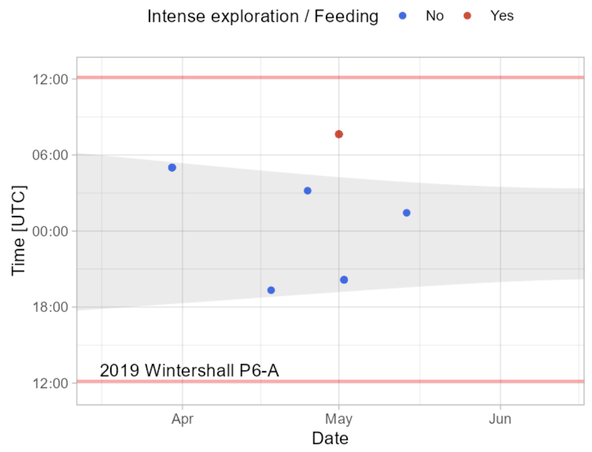


*Figure SF3-33: Recorded acoustic presence of Nathusius' pipistrelle – spring 2019 at Wintershall P6-A*

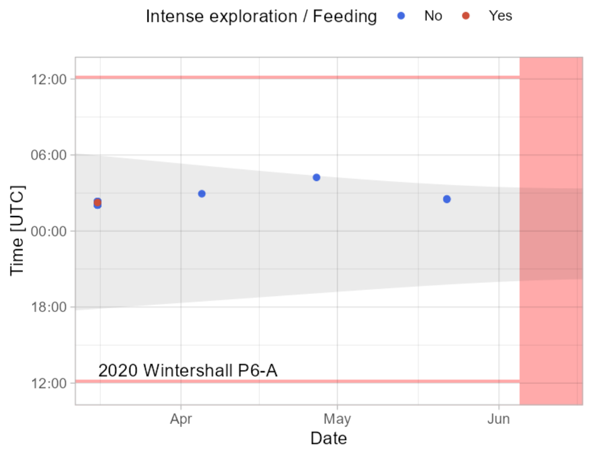


*Figure SF3-34: Recorded acoustic presence of Nathusius' pipistrelle – spring 2020 at Wintershall P6-A*

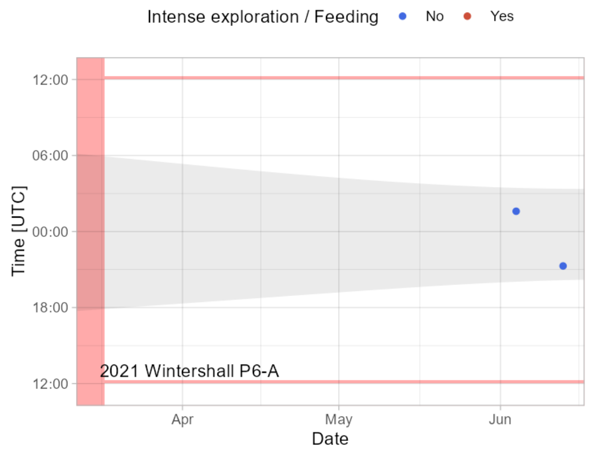


*Figure SF3-35: Recorded acoustic presence of Nathusius' pipistrelle – spring 2021 at Wintershall P6-A*

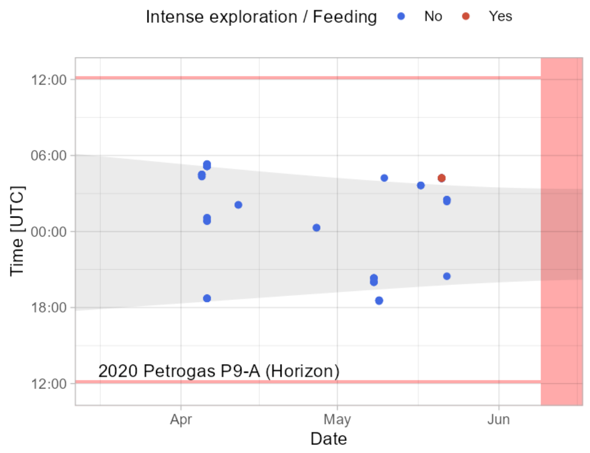


*Figure SF3-36: Recorded acoustic presence of Nathusius' pipistrelle – spring 2019 at Petrogas Q1-A (Helder)*

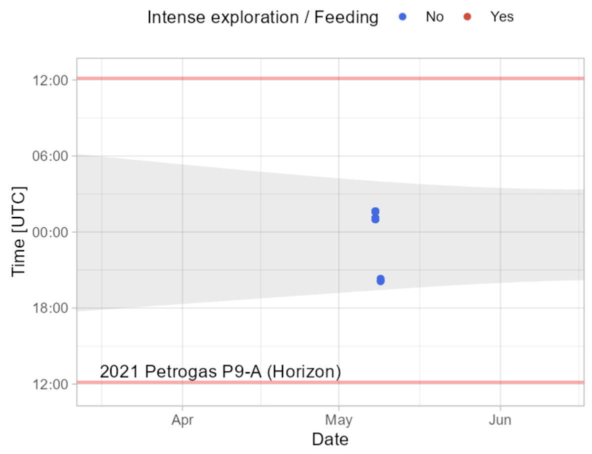


*Figure SF3-37: Recorded acoustic presence of Nathusius' pipistrelle – spring 2020 at Petrogas Q1-A (Helder)*

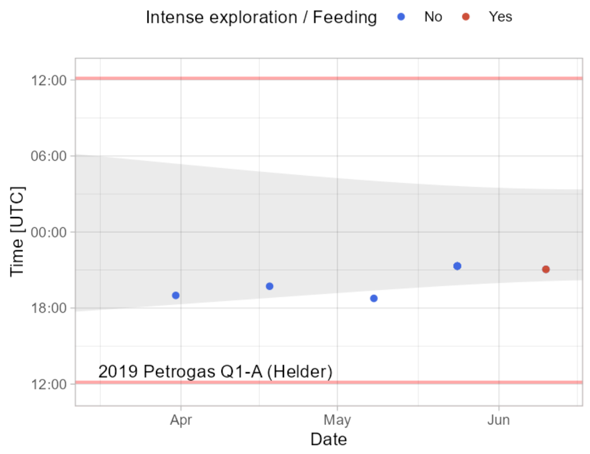


*Figure SF3-38: Recorded acoustic presence of Nathusius' pipistrelle – spring 2019 at Petrogas Q1-A (Helder)*

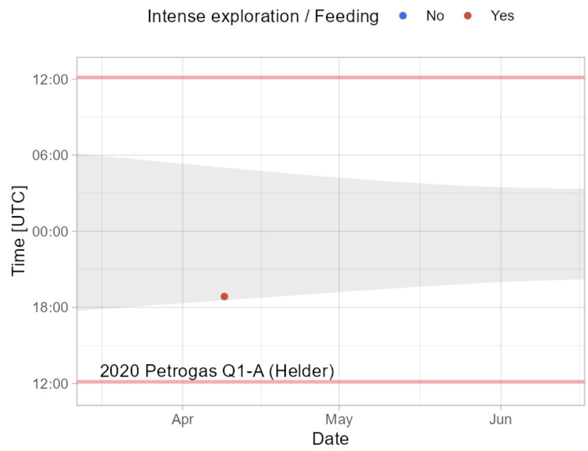


*Figure SF3-39: Recorded acoustic presence of Nathusius' pipistrelle – spring 2020 at Petrogas Q1-A (Helder)*

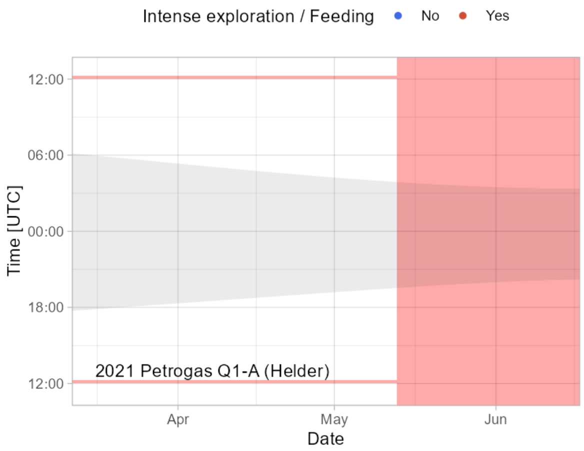


*Figure SF3-40: Recorded acoustic presence of Nathusius' pipistrelle – spring 2021 at Petrogas Q1-A (Helder)*

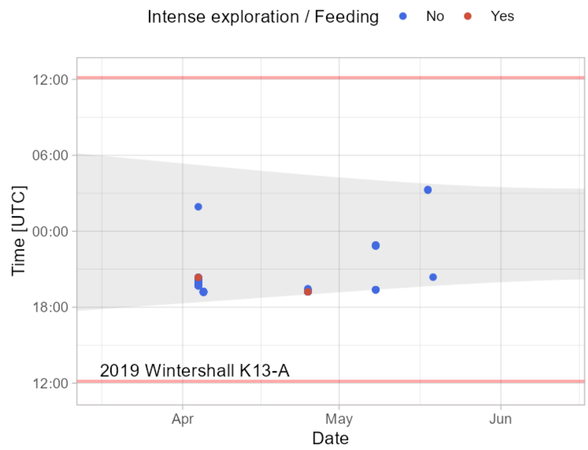


*Figure SF3-41: Recorded acoustic presence of Nathusius' pipistrelle – spring 2019 at Wintershall K13-A*

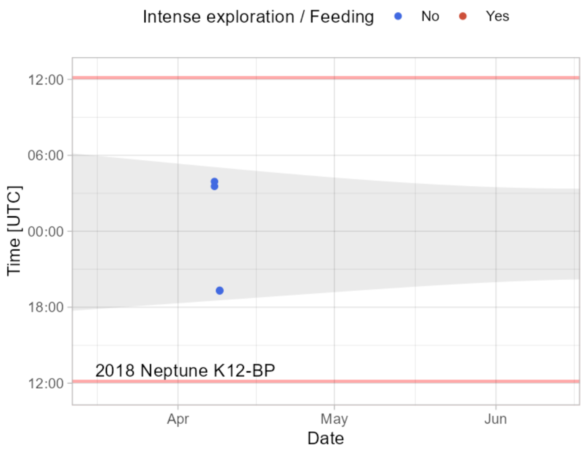


*Figure SF3-42: Recorded acoustic presence of Nathusius' pipistrelle – spring 2018 at Neptune K12-BP*

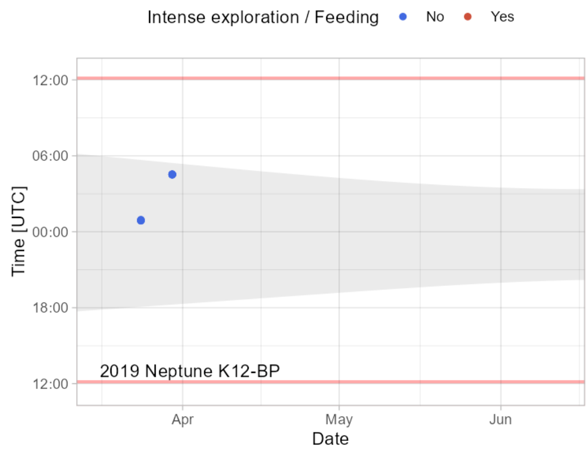


*Figure SF3-43: Recorded acoustic presence of Nathusius' pipistrelle – spring 2019 at Neptune K12-BP*

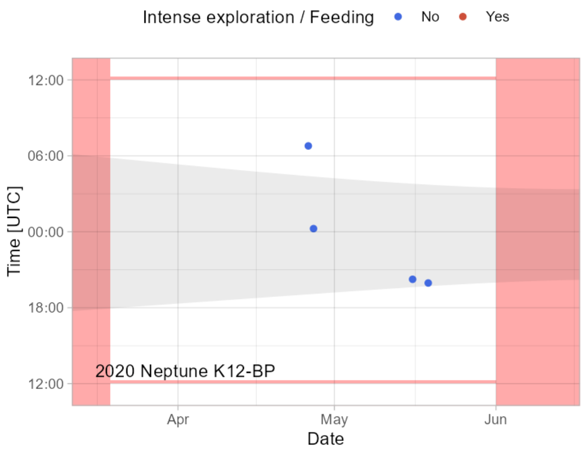


*Figure SF3-44: Recorded acoustic presence of Nathusius' pipistrelle – spring 2020 at Neptune K12-BP*

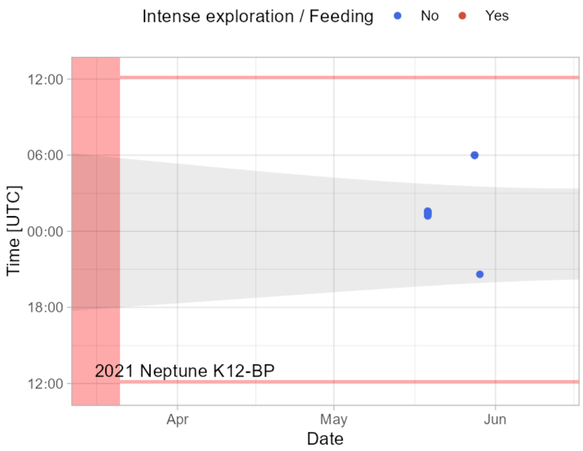


*Figure SF3-45: Recorded acoustic presence of Nathusius' pipistrelle – spring 2021 at Neptune K12-BP*

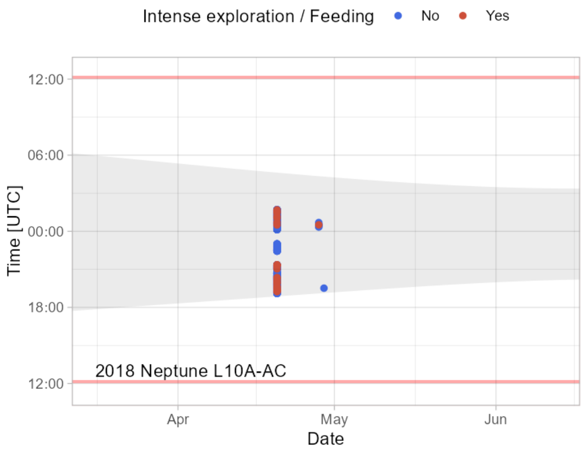


*Figure SF3-46: Recorded acoustic presence of Nathusius' pipistrelle – spring 2018 at Neptune L10A-AC*

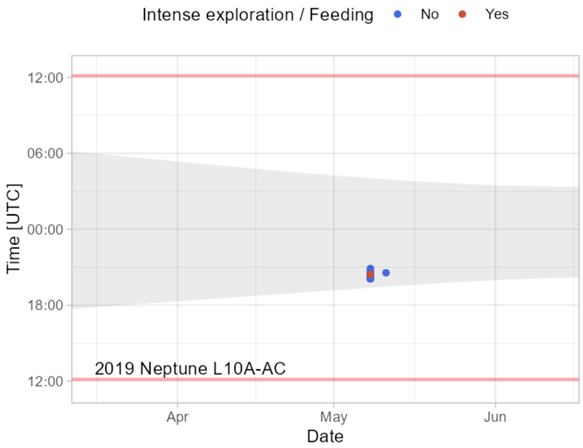


*Figure SF3-47: Recorded acoustic presence of Nathusius' pipistrelle – spring 2019 at Neptune L10A-AC*

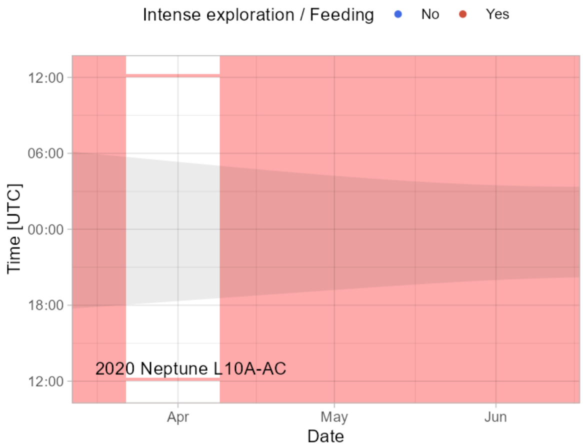


*Figure SF3-48: Recorded acoustic presence of Nathusius' pipistrelle – spring 2020 at Neptune L10A-AC*
