## Supplementary file 4 for "Directional variation and method-specific detection patterns in offshore bat migration: implications for wind farm mitigation"

**Supplementary file 4: tracking and acoustic model output**

<u>Tracking model</u>

The initial fully-parameterized tracking model included the covariates:

- night number (time)
- night number (time) quadratic
- wind speed
- wind speed quadratic
- cos θ
- sin θ
- cos 2θ
- sin 2θ
- wind speed x cos θ
- wind speed x sin θ
- wind speed x cos 2θ
- wind speed x sin 2θ
- sigma individual

Covariates were sequentially removed based on the weakest support (whether the 95% posterior credible interval of their beta parameters overlapped zero). The order in which the covariates were dropped:

1. wind speed x sin θ
2. night number (time) quadratic
3. wind speed x cos 2θ
4. cos 2θ
5. sin θ
6. wind speed x sin 2θ
7. sin 2θ

Convergence was confirmed in the final model for all parameters ($\hat{R} < 1.01$). Table SF1 includes the parameter estimates, Gelman–Rubin convergence diagnostic ($\hat{R}$), effective sample size (n.eff) and the posterior probability the parameter (f). Figure SF1 shows the posterior distributions of the estimated betas.

*Table SF1: tracking model output*

| Parameter | mean | sd | 2.50% | 25% | 50% | 75% | 97.50% | Rhat | n.eff | overlap0 | f |
|---|---|---|---|---|---|---|---|---|---|---|---|
| beta0 | 7.908506 | 3.461004 | 3.095676 | 5.502024 | 7.245227 | 9.586554 | 16.649177 | 1.005886 | 1564 | 0 | 0.999989 |
| beta_time | -0.10353 | 0.072163 | -0.29351 | -0.1357 | -0.08685 | -0.05325 | -0.012592 | 1.006374 | 1598 | 0 | 0.993097 |
| beta_windspeed | -0.26119 | 0.272129 | -0.83509 | -0.43505 | -0.2461 | -0.0721 | 0.2294419 | 1.001039 | 7461 | 1 | 0.833563 |
| beta_windspeed_sq | 0.027186 | 0.014957 | 0.002991 | 0.016196 | 0.025506 | 0.036461 | 0.0606411 | 1.001055 | 7111 | 0 | 0.989615 |
| beta_cos1 | -0.12127 | 0.474875 | -1.05341 | -0.44134 | -0.12106 | 0.199483 | 0.8080528 | 1.000068 | 95972 | 1 | 0.600618 |
| beta_windspeed_cos1 | -0.28899 | 0.123348 | -0.56066 | -0.36421 | -0.27851 | -0.20252 | -0.076828 | 1.001009 | 7260 | 0 | 0.997944 |
| sigma_ind | 1.233672 | 0.877877 | 0.068365 | 0.600998 | 1.074548 | 1.67111 | 3.3967427 | 1.00464 | 1977 | 0 | 1 |
| deviance | 77.23698 | 11.75003 | 53.83714 | 69.27368 | 77.4748 | 85.34155 | 99.633265 | 1.001587 | 4442 | 0 | 1 |

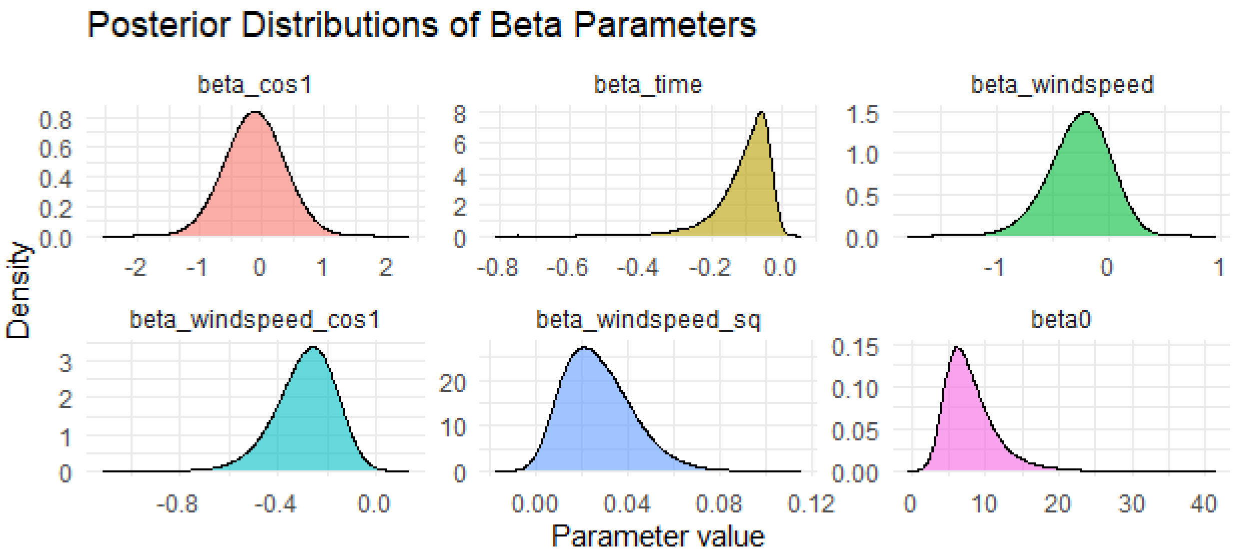


*Figure SF1: posterior distributions of the estimated betas - tracking model.*

<u>Acoustic model</u>

The initial fully-parameterized acoustic model included the covariates:

- night number (time)
- night number (time) quadratic
- wind speed
- wind speed quadratic
- cos θ
- sin θ
- cos 2θ
- sin 2θ
- wind speed x cos θ
- wind speed x sin θ
- wind speed x cos 2θ
- wind speed x sin 2θ
- period
- period x cos θ
- period x sin θ
- period x cos 2θ
- period x sin 2θ
- precipitation
- atmospheric pressure change
- residual temperature on night number (time)
- cloud cover
- lunar phase x cos θ
- lunar phase x sin θ
- lunar phase x cos 2θ
- lunar phase x sin 2θ
- sigma individual

Covariates were sequentially removed based on the weakest support (whether the 95% posterior credible interval of their beta parameters overlapped zero and on the corresponding evidence ratio). The order in which the covariates were dropped:

1. lunar phase x sin 2θ
2. lunar phase x sin θ
3. period x cos θ
4. lunar phase x cos θ
5. lunar phase x cos 2θ
6. cloud cover
7. precipitation
8. period x sin 2θ
9. period x cos 2θ
10. atmospheric pressure change
11. night number (time) quadratic
12. residual temperature on night number (time)
13. wind speed x cos 2θ
14. cos 2θ

Convergence was confirmed in the final model for all parameters ($\hat{R} < 1.01$). Table SF2 includes the parameter estimates, Gelman–Rubin convergence diagnostic ($\hat{R}$), effective sample size (n.eff) and the posterior probability the parameter (f). Figure SF2 shows the posterior distributions of the estimated betas.

*Table SF2: acoustic model output*

| parameter | mean | sd | 2.50% | 25% | 50% | 75% | 97.50% | Rhat | n.eff | overlap0 | f |
|---|---|---|---|---|---|---|---|---|---|---|---|
| beta0 | 7.915514 | 1.320688 | 5.800598 | 6.979426 | 7.75788 | 8.68458 | 10.9106285 | 1.004915 | 2086 | 0 | 1 |
| beta_time | -0.20207 | 0.048745 | -0.312797 | -0.23044 | -0.19624 | -0.16726 | -0.1248131 | 1.005554 | 1829 | 0 | 1 |
| beta_period | 6.272837 | 1.668535 | 3.629972 | 5.082138 | 6.071495 | 7.237223 | 10.0798604 | 1.00533 | 1875 | 0 | 1 |
| beta_windspeed | -0.00558 | 0.085554 | -0.179137 | -0.06214 | -0.00336 | 0.053146 | 0.15614617 | 1.000292 | 24327 | 1 | 0.515823 |
| beta_windspeed_sq | 0.011136 | 0.00517 | 0.001587 | 0.007543 | 0.010928 | 0.014509 | 0.02182917 | 1.000367 | 18703 | 0 | 0.990168 |
| beta_cos1 | 0.405202 | 0.312451 | -0.2074 | 0.195046 | 0.405007 | 0.615116 | 1.0188709 | 1.000064 | 104926 | 1 | 0.903169 |
| beta_sin1 | -0.45249 | 0.415183 | -1.280823 | -0.72889 | -0.44766 | -0.17072 | 0.34926753 | 1.000097 | 76622 | 1 | 0.863472 |
| beta_sin2 | -0.70931 | 0.330528 | -1.356355 | -0.93125 | -0.7102 | -0.48785 | -0.0576457 | 1.000069 | 104508 | 0 | 0.983371 |
| beta_windspeed_cos1 | -0.16542 | 0.048392 | -0.26298 | -0.1974 | -0.16459 | -0.1324 | -0.0729943 | 1.00017 | 39532 | 0 | 0.999832 |
| beta_windspeed_sin1 | -0.20913 | 0.052289 | -0.314409 | -0.24372 | -0.20816 | -0.17349 | -0.1092526 | 1.000241 | 27969 | 0 | 0.999994 |
| beta_windspeed_sin2 | 0.157952 | 0.045772 | 0.068905 | 0.127031 | 0.157629 | 0.188542 | 0.24861888 | 1.000092 | 76624 | 0 | 0.999759 |
| beta_sin1_period | 1.437895 | 0.38167 | 0.703627 | 1.179013 | 1.432165 | 1.690717 | 2.20316383 | 1.00007 | 97788 | 0 | 0.999952 |
| sigma_ind | 1.574849 | 0.548115 | 0.672951 | 1.190297 | 1.516872 | 1.895804 | 2.80418905 | 1.004859 | 2012 | 0 | 1 |
| deviance | 870.3318 | 55.22984 | 756.3681 | 833.9633 | 872.7028 | 909.3069 | 970.85913 | 1.003421 | 2374 | 0 | 1 |

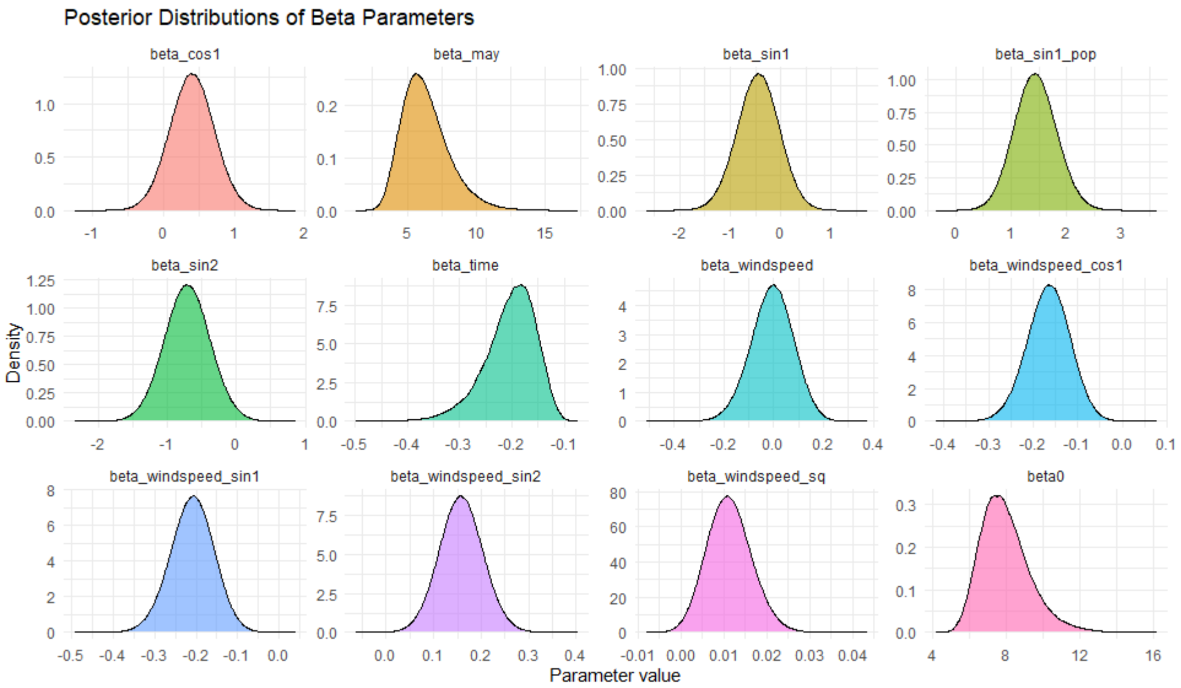


*Figure SF2: posterior distributions of the estimated betas – acoustic model.*
